# Beyond Glycogen Storage: AMPKγ2 Regulates Cardiac Hypertrophy and Electrophysiology via Myosin Interaction

**DOI:** 10.64898/2026.04.20.719766

**Authors:** Qianyun Ge, Kusumika Saha, Micah L. Burch, Willow H. Battista, KC Ashmita, Max Homilius, Rachelle A. Victorio, Dajun Quan, Hsiang-Ling Huang, Joseph M. Hazel, Alex Williams, Emily Pan, Krishna Chinthalapudi, Sarah M. Heissler, Calum A. MacRae, Wandi Zhu

**Affiliations:** Department of Molecular Medicine and Therapeutics, The Ohio State University, College of Medicine, Columbus, 43210, USA; Department of Medicine, Brigham and Women’s Hospital, Boston, MA 02115, USA; Harvard Medical School, Boston, MA 02115, USA; Department of Physiology and Cell Biology, The Ohio State University, College of Medicine, Columbus, 43210, USA; Dorothy M. Davis Heart and Lung Research Institute, The Ohio State University, College of Medicine, Columbus, 43210, USA

## Abstract

**Introduction:** Variants in *PRKAG2* cause hypertrophic cardiomyopathy (HCM) and conduction disturbances. While prior studies associated *PRKAG2*-related hypertrophy with increased glycogen storage, many HCM phenotypes remain unexplained. We aimed to uncover how *PRKAG2* variants induce myocyte hypertrophy and electrical changes during early cardiac development.

**Methods:** We generated transgenic zebrafish expressing wild-type (Tg^WT^) or pathogenic variant (Tg^R299Q^) *Prkag2* cDNA under a myocardium-specific promoter, and examined cardiac electrophysiology, contractile function, and cytoarchitecture during cardiogenesis and in adult hearts.

**Results:** Tg^R299Q^ fish showed hypertrophic cardiomyocytes and progressive contractile abnormalities, recapitulating human HCM phenotypes. Cardiomyocyte glycogen was elevated in adult but not embryonic hearts. Despite the absence of glycogen accumulation at 6-day post-fertilization, Tg^R299Q^ hearts showed electrical abnormalities, including reduced conduction velocity and prolonged action potential and Ca^2+^ transient durations. We observed decreased AMPK phosphorylation in the Tg^R299Q^ hearts. However, AMPK activation did not rescue the electrophysiological abnormalities in Tg^R299Q^. Proximity ligation assays and co-immunoprecipitation identified a physical interaction between AMPKγ2 and myosin, enhanced by the R299Q variant and accompanied by increased AMPKγ2 localization to the myofilament. Na⁺/Ca²⁺ exchanger (NCX) inhibition increased Ca^2+^ duration and diastolic Ca^2+^ in Tg^WT^ but not Tg^R299Q^ hearts, indicating reduced free cytosolic Ca^2+^ for NCX-mediated extrusion in Tg^R299Q^. These findings suggest that enhanced AMPKγ2-myosin interaction may promote myofilament Ca²⁺ retention, thereby prolonging Ca²⁺ transient duration and APD in the mutant. Notably, the myosin inhibitor mavacamten reduced AMPKγ2-myosin interaction in Tg^R299Q^ hearts, and both mavacamten and *vmhcl* knockdown rescued the early electrophysiological abnormalities.

**Conclusions:** The *PRKAG2* variant altered cardiac excitability, contractility, and Ca^2+^ handling during cardiogenesis, independent of glycogen accumulation. Enhanced interactions between AMPKγ2 and myosin contributed to these early changes. Our study revealed a novel link between cellular energy sensing and contractile machinery, with therapeutic potential for modulating contractile function in cardiomyopathies.

## Introduction

Hypertrophic cardiomyopathy (HCM) is the most prevalent form of Mendelian-inherited heart disease, characterized by abnormal thickening of the heart muscle^1^. While the majority of HCM cases can be attributed to variants in genes encoding sarcomere proteins, genetic variants in cell metabolic regulators can also lead to clinical HCM phenotypes^1,2^. One such example is the *PRKAG2* gene, which encodes the γ2 subunit of the AMP-activated protein kinase (AMPK). The γ subunits of AMPK play a crucial role as energy sensors within cells by competitively binding to adenyl nucleotides^3^. Each γ subunit contains four tandem cystathionine β-synthase (CBS) domains that can bind to AMPs, ATPs, and ADPs^3,4^. In response to cellular energy stress, reflected as elevated AMP/ATP and ADP/ATP ratios within the cell, the γ subunit undergoes differential binding to these nucleotides. Subsequently, the γ subunit phosphorylates and activates AMPK, promoting increased catabolism to restore energy homeostasis^3^.

Variants in *PRKAG2* affecting the function of the γ2 subunit lead to heterogeneous clinical phenotypes, including early onset HCM and progressive conduction disorders such as ventricular pre-excitation and atrial arrhythmias^5–8^. While both sarcomere gene-linked and *PRKAG2*-linked HCMs exhibit similar left ventricular hypertrophy phenotypes, variants in *PRKAG2* are associated with more electrical disturbances and an increased rate of progression to heart failure and increased incidence of sudden cardiac death in affected individuals^8–10^.

Previous studies using transgenic mouse and human induced pluripotent stem cell-derived cardiomyocyte models of *PRKAG2* variants have revealed elevated glycogen stores and the presence of glycogen-containing vacuoles in cardiomyocytes^11–13^. This increased glycogen accumulation has been thought to contribute to the pathogenesis of HCM and conduction disturbances^11–13^. However, many of the observed HCM phenotypes in *PRKAG2* disorder, including changes in myocyte contractility and cell excitability, cannot be solely explained by the mass effect of glycogen and secondary changes in overall cell size. In fact, it has been demonstrated that modulating glycogen synthase (GYS1) to reduce glycogen storage in the *PRKAG2* genetic mouse model was not sufficient to reduce cardiac hypertrophy^14^, suggesting that additional mechanisms underlie *PRKAG2*-linked HCM. Furthermore, prior investigations have primarily focused on examining mature cardiac tissue, with limited information regarding how *PRKAG2* regulates early heart development and cardiomyocyte cytoarchitecture and functional establishment during embryogenesis.

To address these gaps, we developed transgenic zebrafish models of the *PRKAG2* syndrome to investigate its role in regulating cardiac function and cardiomyocyte morphology during early cardiogenesis and in mature hearts. We used a combination of genetics, electrophysiology, biochemistry, cell biology, and transcriptomic methods to uncover the molecular mechanisms by which a *PRKAG2* variant alters cardiomyocyte excitability, Ca^2+^ handling, and the regulation of myofilament proteins in early heart development, preceding glycogen accumulation in cardiomyocytes. Our study not only discovered early pathophysiologic effects of mutant *PRKAG2* independent of glycogen storage but also revealed shared molecular mechanisms between sarcomere-linked and *PRKAG2*-associated HCMs, suggesting alternative therapeutic approaches for the treatment of multiple forms of HCM and glycogen storage myopathies.

## Results

### Transgenic expression of the *Prkag2* R299Q variant in zebrafish causes cardiomyocyte hypertrophy

To model *PRKAG2* disorder in zebrafish, we generated two transgenic models: one expressing the cDNA of murine wild-type *Prkag2* (Tg^WT^) and another expressing *Prkag2* carrying the R299Q point mutation (Tg^R299Q^), under a myocardial-specific *cmlc2* promoter. The R299Q variant in murine *Prkag2* is equivalent to the R302Q variant in the human isoform, which is one of the most common pathological *PRKAG2* variants found in patients with HCM^10^.

At the adult stage (6-12 months post-fertilization), atrial and ventricular dimensions were substantially enlarged in the Tg^R299Q^ fish, compared to the *wild-type* (TuAB) and Tg^WT^ (**Figure 1A**), mirroring human PRKAG2 syndrome characteristics of atrial enlargement and ventricular hypertrophy^7^. To assess ventricular cardiomyocyte sizes, we used both histological staining and cell capacitance measurements through patch clamp recordings. Wheat germ agglutinin (WGA) stain of longitudinal sections of the ventricles clearly delineated cell borders (**Figure 1B**). Quantitative analysis of cell size and shape from ventricular section images revealed elevated cardiomyocyte area and cell width in Tg^R299Q^, compared to Tg^WT^ and TuAB (**Figure 1C, 1D**). There were no significant differences in cardiomyocyte length between Tg^WT^ and Tg^R299Q^ (**Figure 1E**). Measurement of cell membrane capacitance of isolated cardiomyocytes using patch clamp showed an increased capacitance in ventricular myocytes of Tg^R299Q^ compared to TuAB, suggesting an augmented cell surface area in Tg^R299Q^ (**Supplement Figure 1A**). No difference in membrane capacitance of atrial cardiomyocytes was observed between the TuAB and Tg^R299Q^ (**Supplement Figure 1B**). We also examined the total cardiomyocyte number among the three genotypes at 3-day post-fertilization (dpf), when the heart just completes looping. Both the Tg^WT^ and Tg^R299Q^ embryonic hearts showed an increase in cardiomyocyte counts in atria and ventricles, compared to TuAB (**Supplement Figure 1C-E**). However, there was no significant difference between Tg^WT^ and Tg^R299Q^ (**Supplement Figure 1C-E**). These findings suggest that the R299Q variant in *Prkag2* leads to a ventricular myocyte hypertrophy phenotype in zebrafish, primarily characterized by an increase in cell width. Atrial enlargement in Tg^R299Q^ fish is likely a primary phenotype, given that atrial abnormalities frequently occur independently of ventricular hypertrophy in human *PRKAG2* syndrome^7,15^, although a secondary contribution from ventricular dysfunction may also play a role.

**Figure 1.**
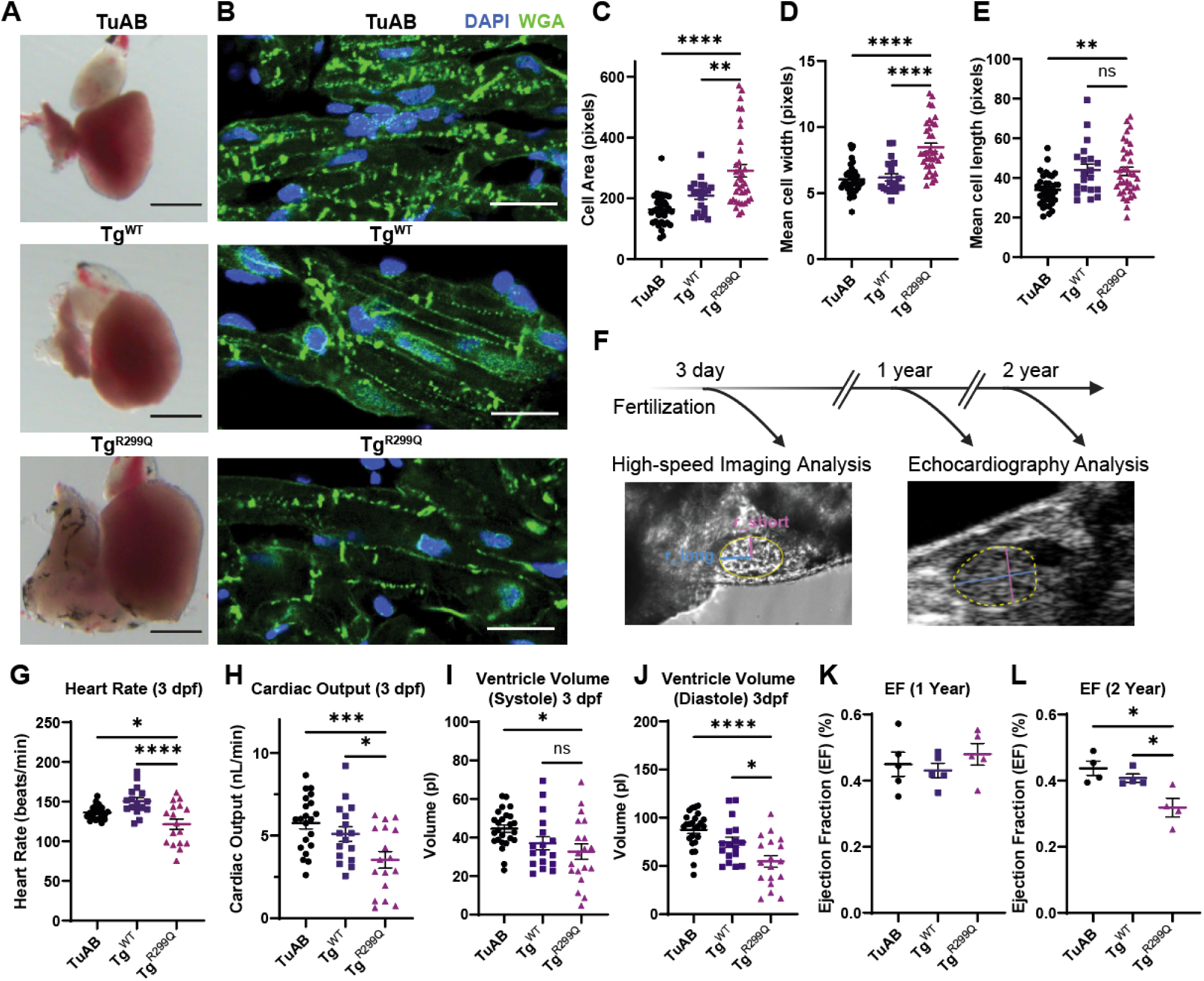
Transgenic expression of the pathological *Prkag2* variant R299Q induces ventricular cardiomyocyte hypertrophy. A. Brightfield images showing the morphology of hearts dissected from adult TuAB, Tg^WT^, and Tg^R299Q^ fish. B. Confocal images of ventricular cryosections of TuAB, Tg^WT^, and Tg^R299Q^ adult fish hearts stained with WGA for cell border and DAPI for nuclei. C-E. Quantification of cell area, width, and length from confocal images of longitudinal ventricular sections stained with DAPI and WGA. Each data point represents a cell, and four adult hearts from each genotype were used for the analysis. F. Cardiac functions of different genotypes were studied using high-speed imaging at the embryonic stage (3 dpf) and echocardiography at the adult stage (1 or 2 years post-fertilization). Exemplary images showing the ventricle contour and short and long axes of the ventricle. G-J. Quantified heart rate (G), cardiac output (H), ventricular systolic (I), and diastolic volume (J) of 3 dpf TuAB, Tg^WT^, and Tg^R299Q^ embryos. Each data point represents an individual embryo. K-L. Ejection fraction (EF) quantified from the echocardiography analyses of adult TuAB, Tg^WT^, and Tg^R299Q^ fish at 1 year (K) and 2 years (L) post-fertilization. Each data point represents an individual animal. Results are expressed as mean ± S.E.M. *P < 0.05, **P < 0.01, ***P<0.001, and ****P < 0.0001.

### Tg^R299Q^ heart exhibited progressive abnormalities in cardiac contractile function

We next characterized the effect of the *Prkag2* R299Q variant on heart function during early cardiogenesis (3 dpf) and through aging (1 or 2-year post-fertilization) using high-speed imaging and echocardiography analyses (**Figure 1F**). At 3 dpf, Tg^R299Q^ exhibited a decreased heart rate compared to Tg^WT^ and TuAB (**Figure 1G**). This finding is consistent with prior studies in murine models and humans showing AMPKγ2 as an energy sensor affecting intrinsic heart rate^16,17^. Additionally, we observed a decrease in ventricular cardiac output, with an unchanged systolic volume, but reduced diastolic volume in 3 dpf Tg^R299Q^ compared to Tg^WT^ and TuAB (**Figure 1H-J**), suggesting R299Q already impaired relaxation during early cardiogenesis. In mature hearts, at 1-year post-fertilization, there was no difference in ventricular ejection fraction among the TuAB, Tg^WT^, and Tg^R299Q^ (**Figure 1K**), similar to human clinical echocardiography findings^16^. However, the Transmission Electron Microscopy (TEM) images of ventricular longitudinal sections revealed a significant reduction in sarcomere length in Tg^R299Q^ myocytes compared to TuAB and Tg^WT^, suggesting that the *Prkag2* variant affects sarcomere organization and contractile function (**Supplement Figure 2**). In aged zebrafish (2-year-old), Tg^R299Q^ fish showed a reduced ejection fraction compared to Tg^WT^ and TuAB (**Figure 1L**), a characteristic of heart failure. Our results showed a progressive change in heart contractile function in the Tg^R299Q^ model, with abnormalities observed as early as 3 dpf.

### Glycogen storage was unaffected by the R299Q variant during early cardiogenesis

Excess glycogen deposition has been implicated in the pathogenesis of *PRKAG2* disorder^18^. Given the progressive nature of glycogen accumulation in cells, we sought to examine the glycogen content in hearts at both adult and early embryogenesis stages. Using a glycogen assay, we first quantified the absolute glycogen content in lysates of the excised hearts from adult and 6 dpf embryonic fish. In adult hearts, the glycogen content was found to be increased by threefold in Tg^R299Q^, compared to TuAB and Tg^WT^ (**Figure 2A**). In contrast, there was no discernible difference in glycogen content among the three genotypes at 6 dpf (**Figure 2B**). Consistent with the results from the glycogen assay, the Periodic Acid-Schiff (PAS) stain of 6 dpf embryonic hearts showed no difference in glycogen stores between TuAB and Tg^R299Q^ (**Figure 2C**). These results suggest that while glycogen stores were upregulated in mature heart samples of Tg^R299Q^, there was no evidence of glycogen accumulation during early cardiogenesis.

**Figure 2.**
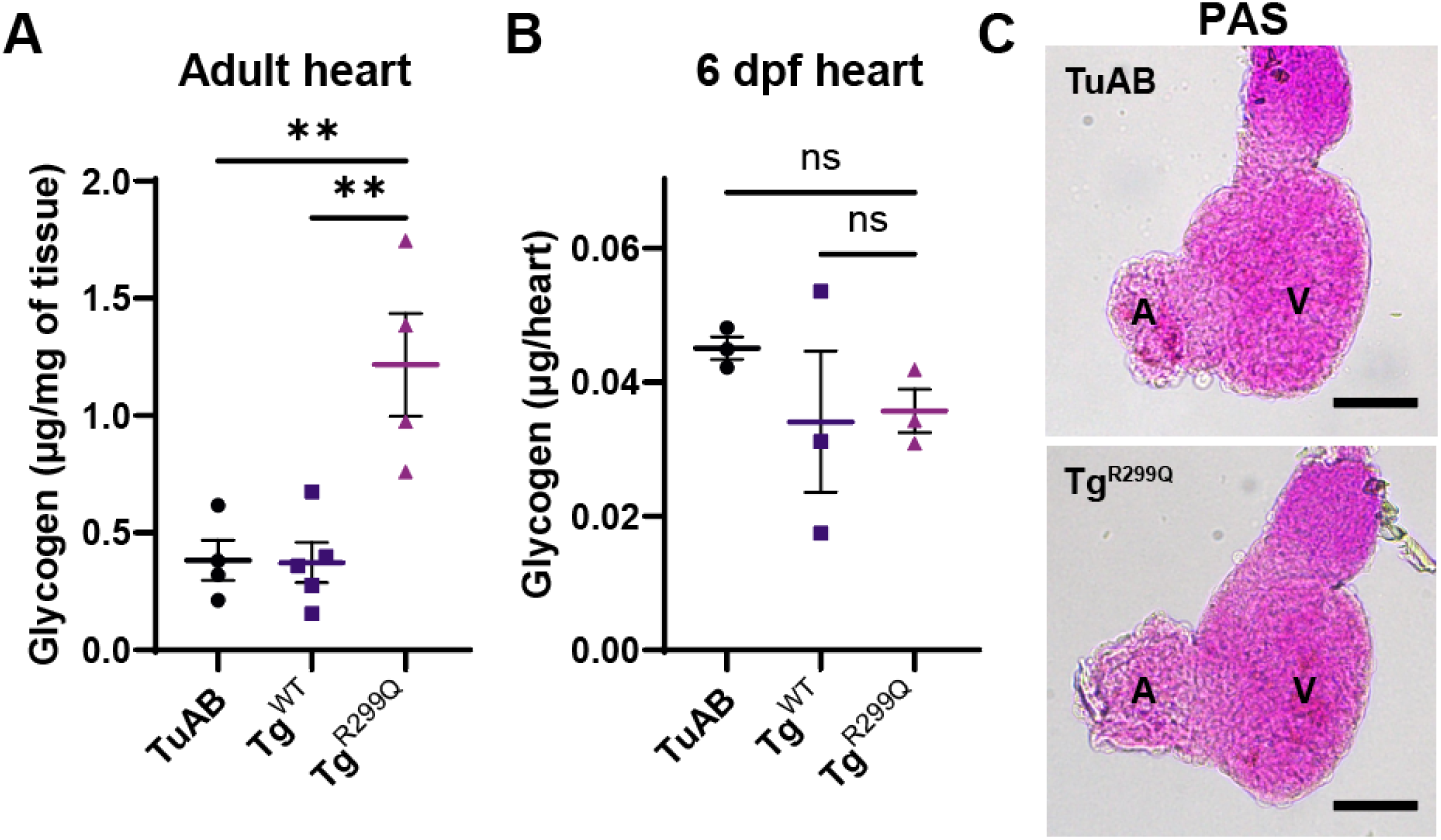
Tg^R299Q^ exhibited an increase in glycogen storage in adult hearts, but not in embryonic hearts. A. Glycogen content from adult heart lysates, which was determined with a fluorometric glycogen assay kit. Each point represents the mean of three technical replicates of one heart sample. B. Glycogen content that was measured with a fluorometric glycogen assay from 6 dpf embryonic heart lysates. Each lysate sample contains 10 isolated embryonic hearts. Each point represents the mean of three technical replicates of one lysate sample. C. Representative images of periodic acid–Schiff (PAS) stain of fixed embryonic hearts from 6 dpf TuAB and Tg^R299Q^ larvae. Results are expressed as mean ± S.E.M. **P < 0.01.

### The *Prkag2* variant caused electrical and Ca^2+^ handling abnormalities in developing hearts

Since *PRKAG2* variants commonly manifest with abnormalities of cardiac electrical conduction and excitability in clinical presentations, we aimed to test whether these electrical disturbances arise during early cardiogenesis despite the absence of glycogen storage abnormality. Using high-speed optical mapping of transmembrane potential with the voltage-sensitive dye FluoVolt, we were able to capture cardiac conduction at cellular resolution in 6 dpf embryonic hearts. We found that the conduction velocity of electrical impulses was significantly slowed in the atrium and ventricle of Tg^R299Q^, compared to TuAB and Tg^WT^ (**Figure 3A-C**). The maximum upstroke velocity of action potentials (Vmax) in atria and ventricles remained largely unchanged between Tg^WT^ and Tg^R299Q^ (**Supplement Figure 3A-B**). The reduced conduction associated with the R299Q variant suggests a decrease in cardiomyocyte electrical coupling or a reduction in the availability of voltage-gated Na^+^ channels in the *Prkag2* mutant.

**Figure 3.**
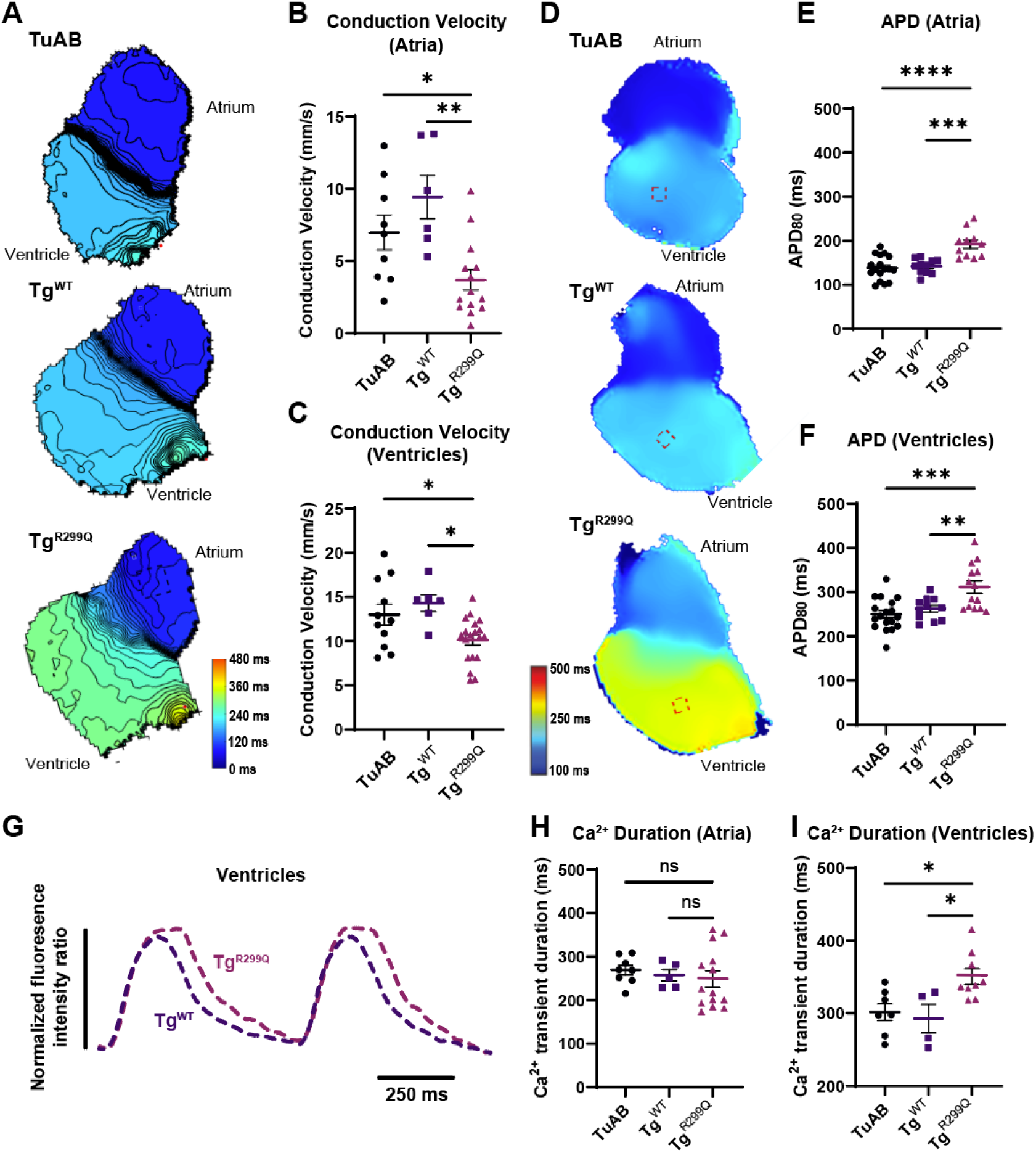
Tg^R299Q^ embryonic hearts exhibited abnormal electrical activities and Ca^2+^ handling. A. Representative isochrone maps of hearts isolated from 6 dpf TuAB, Tg^WT^, and Tg^R299Q^ zebrafish larvae optically mapped with a voltage-sensitive dye FluoVolt. The color scale depicts the timing of electrical activation. B-C. Calculated atrial and ventricular conduction velocities from 6 dpf TuAB, Tg^WT^, and Tg^R299Q^ hearts. D. Representative maps of action potential duration at 80% repolarization (APD_80_) in isolated hearts field stimulated at 80 beats per minute (750 ms cycle length). The dotted squares reflect the ventricular areas where APD_80_ was measured. E-F. Calculated APD_80_ from atria (E) and ventricles (F) of 6 dpf TuAB, Tg^WT^, and Tg^R299Q^ hearts. G. Representative normalized ventricular Ca^2+^ transient measured by Fura 2 comparing Tg^WT^ and Tg^R299Q^. H-I. Duration of the Ca^2+^ transient (at 70% decay) in atria (H) and ventricles (I) of 6 dpf TuAB, Tg^WT^, and Tg^R299Q^ hearts. [Ca^2+^]_i_ was calculated by fluorescence ratio (F340/F380). Results are expressed as mean ± S.E.M. Each data point represents the mean of an individual heart studied. *P < 0.05, **P < 0.01, ***P < 0.001, and ****P < 0.0001.

We also measured the action potential duration (APD) in the atria and ventricles with a controlled field electrical stimulation at a cycle length of 750 ms (**Figure 4F**). Compared to TuAB and Tg^WT^, Tg^R299Q^ exhibited prolonged APD_80_ (time to 80% recovery from peak to resting potential) in both atria and ventricles (**Figure 3D-F**). The prolongation of APD could be attributed to an increase in persistent Na^+^ current, extended Ca^2+^ transient duration, or a reduction in repolarizing K^+^ current. Notably, RNA sequencing results revealed suppressed AP repolarization processes (**Supplement Figure 4A**), particularly a decrease in the expression of transcripts of *kcnh6a* (**Supplement Figure 4B**), encoding the zebrafish homology of the hERG channel, which forms a major repolarizing current I_Kr_.

**Figure 4.**
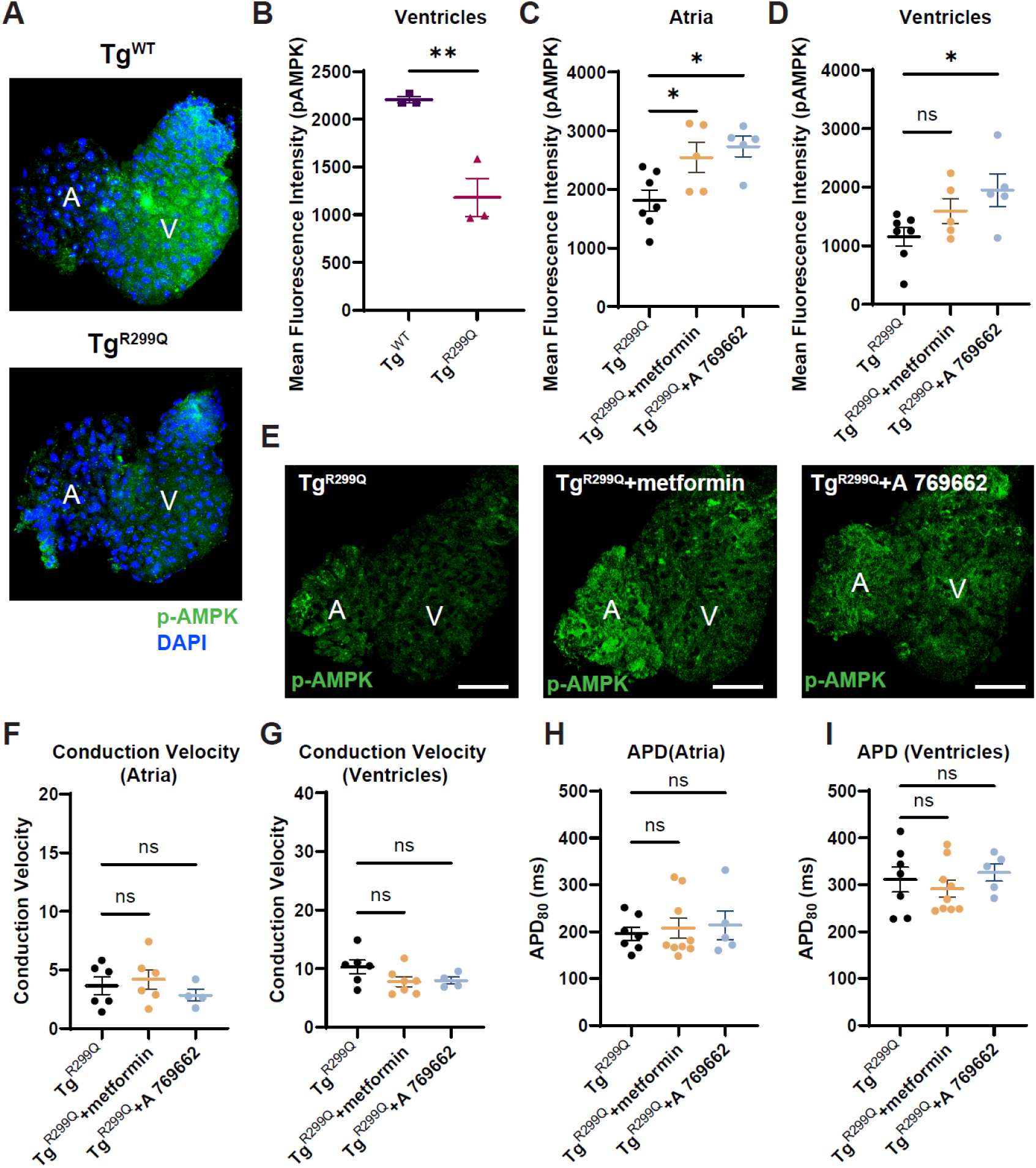
Electrophysiological abnormalities in Tg^R299Q^ hearts are not rescued by modulation of AMPK catalytic activity. A. Representative confocal images of 6 dpf embryonic hearts immuno-stained with the phosphorylated AMPK (Thr172) antibody and DAPI, comparing Tg^WT^ and Tg^R299Q^. B. Quantified mean fluorescence intensity of the phospho-AMPK (p-AMPK) stained ventricles from Tg^WT^ and Tg^R299Q^ 6 dpf hearts. C-D. Quantified mean fluorescence intensity of the p-AMPK stained atria (C) and ventricles (D) from Tg^R299Q^ control (treated with DMSO), Tg^R299Q^ treated with 50 µM metformin, and Tg^R299Q^ treated with 10 µM A 769662 6 dpf hearts. E. Representative confocal images of 6 dpf embryonic hearts immuno-stained with the phosphorylated AMPK (Thr^172^) antibody, comparing Tg^R299Q^ control, treated with metformin, and A 769662. F-G. Conduction velocity of 6 dpf hearts of Tg^R299Q^ control, treated with metformin and A 769662 in atria (F) and ventricles (G). H-I. Measured AP duration (80%) of atria (H) and ventricles (I) from 6 dpf hearts of Tg^R299Q^ control, treated with metformin and A 769662. Results are expressed as mean ± S.E.M. Each data point represents the mean of an individual heart studied. *P < 0.05, and **P < 0.01.

Sarcomere gene-linked HCM is known to alter Ca^2+^ transient in cardiomyocytes due to slowed Ca^2+^ dissociation from myofilaments and limited energy supply for Ca^2+^ uptake and extrusion by Sarcoplasmic Reticulum Ca^2+^ ATPase and Na^+^/Ca^2+^ exchanger (NCX)^1^. However, the impact of *PRKAG2* variants on cardiomyocyte Ca^2+^ handling remains less clear. To assess Ca^2+^ dynamics, we applied high-speed ratiometric Ca^2+^ imaging on isolated 6 dpf embryonic hearts. Our analyses revealed a prolonged Ca^2+^ transient duration in Tg^R299Q^ ventricles, compared to Tg^WT^ and TuAB (**Figure 3G, I**). The atrial Ca^2+^ transient duration remains unaffected in Tg^R299Q^ (**Figure 3H**). We observed no significant difference in diastolic [Ca^2+^]_i_ or the amplitude of Ca^2+^ transient in atria and ventricles between Tg^WT^ and Tg^R299Q^, although both transgenic models showed elevated diastolic [Ca^2+^]_i_ and decreased Ca^2+^ transient amplitude when compared to TuAB (**Supplement Figure 3C-F**). These results suggest that transgenic expression of AMPKγ2 can modulate myocyte Ca^2+^ homeostasis, and the R299Q variant primarily extends the Ca^2+^ transient duration in the ventricle, which can contribute to the prolongation of APD. We found that the Tg^R299Q^ exhibited electrical and Ca^2+^ handling abnormalities during early cardiogenesis, preceding glycogen accumulation, suggesting alternative mechanisms underlying these changes.

### Modulating AMPK activity did not rescue *PRKAG2*-linked electrical abnormalities

As the R299Q variant resides in the CBS1 domain of the γ2 subunit, which can affect adenyl nucleotide binding and potentially alter AMPK activation^4^, we next sought to investigate how AMPK α subunit phosphorylation is altered in the transgenic fish models. In 6 dpf hearts, immunostaining results showed a decreased level of phosphorylation of threonine 172 (Thr172) of AMPKα in Tg^R299Q^, compared to Tg^WT^ (**Figure 4A, B**), suggesting reduced AMPK activation. In adult hearts, western blot analyses showed elevated levels of total AMPKα (**Supplement Figure 5A, B**), but a reduced ratio of phosphorylated AMPKα to total AMPKα in Tg^R299Q^, compared to TuAB, but not compared to Tg^WT^ (**Supplement Figure 5C, D**). These results showed that the R299Q variant reduced AMPK activation in early cardiogenesis, with a lesser degree of inhibition of AMPK activation in mature hearts.

To test whether the electrophysiological changes caused by R299Q are the consequence of a reduced AMPK activation, we treated the Tg^R299Q^ 2 dpf embryos with two different AMPK activators, metformin and A 769662, and assessed cardiac electrophysiological responses with optical mapping at 6 dpf. We found that both compounds effectively increased AMPKα phosphorylation in the atria of the Tg^R299Q^ 6 dpf hearts compared to the untreated condition (**Figure 4C, D, E**). A 769662 also elevated phosphorylated AMPKα levels in the ventricles of Tg^R299Q^ compared to untreated (**Figure 4C**). However, treatment with neither compound was effective in restoring the conduction velocity (**Figure 4F, G)** or APD_80_ (**Figure 4H, I)** changes observed in Tg^R299Q^, suggesting the electrical phenotypes in Tg^R299Q^ developing hearts were not due to alterations in AMPK activation.

### The R299Q variant modulated the subcellular localization of the γ2 subunit and altered myocyte cytoarchitecture

Given the observed early electrophysiology phenotypes and altered AMPK phosphorylation in Tg^R299Q^ at early embryogenesis, we next sought to investigate the impact of this variant on the expression and subcellular localization of the γ2 subunit in embryonic hearts. Immunofluorescence staining of 3 dpf ventricles revealed that the γ2 subunit in TuAB and Tg^WT^ exhibited a cytosolic localization pattern, resembling that of mitochondria (**Figure 5A**). Remarkably, in Tg^R299Q^ ventricular myocytes, AMPKγ2 was redistributed toward the cell periphery (**Figure 5A**). Co-immunostaining with the sarcomere protein Tropomyosin 1 (TPM1) (**Figure 5B**) and myosin heavy chain (MYH; detected with the F59 antibody, which recognizes multiple MYH isoforms, hereafter referred to as myosin) (**Figure 5C**) demonstrated co-localization of AMPKγ2 carrying the R299Q mutation with the sarcomere structure. However, this distinct expression pattern of the mutant γ2 subunit did not impact the localization of the phosphorylated AMPK α subunit (**Figure 5D**).

**Figure 5.**
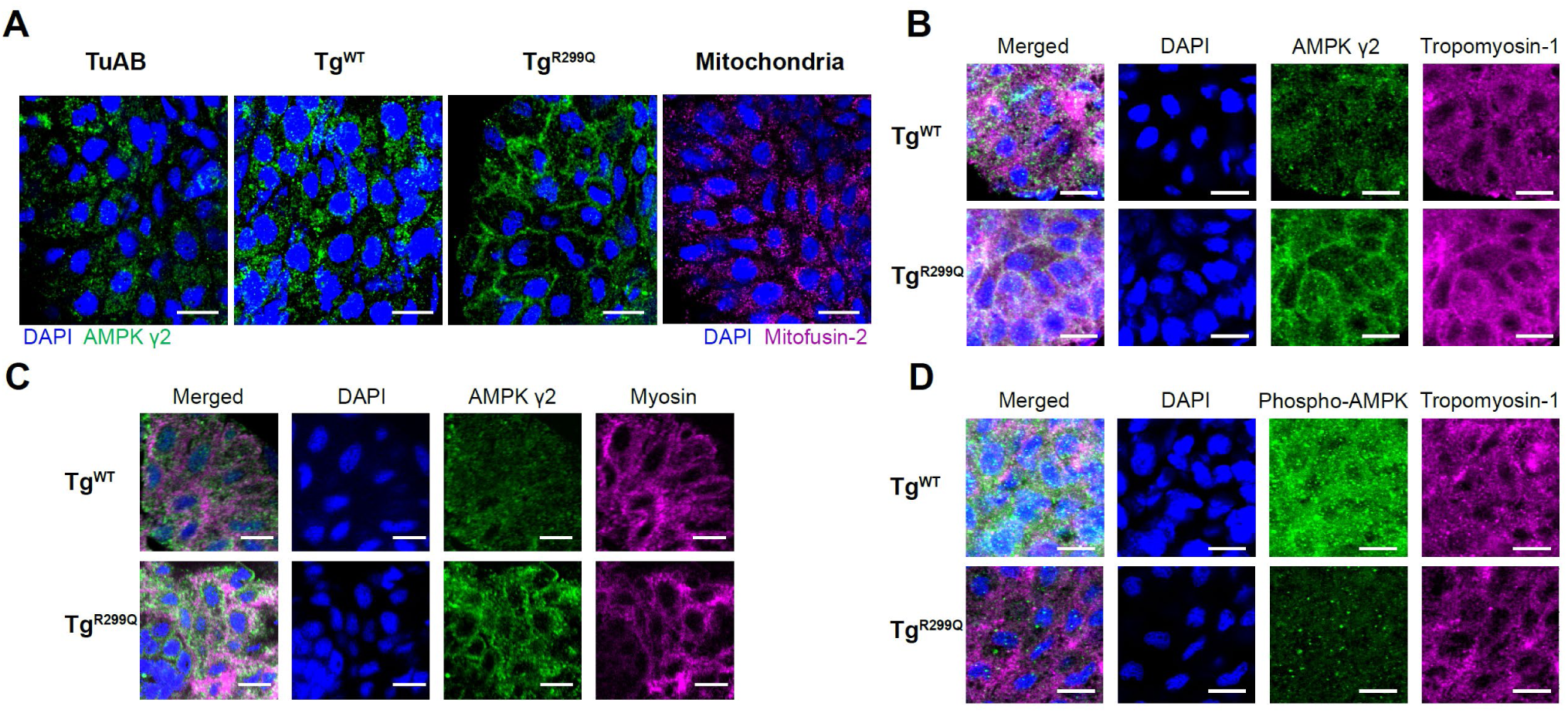
The R299Q variant translocases the AMPKγ2 to the sarcomere. A. Representative confocal images of immunofluorescence labeling of AMPKγ2 and DAPI nuclear stain in the ventricles of TuAB, Tg^WT^, and Tg^R299Q^ 6dpf hearts (left), and images of the ventricle of 3 dpf TuAB heart stained with Mitofusin-2 antibody and DAPI. The scale bars shown are 10µm. B-D. Example immunofluorescence images of ventricles from Tg^WT^ and Tg^R299Q^ 3 dpf fish, stained with AMPKγ2 (green), Tropomyosin-1 (magenta), and DAPI (blue) in (B), AMPKγ2 (green), MYH (magenta), and DAPI (blue) in (C). Phospho-AMPK (green), Tropomyosin-1 (magenta), and DAPI (blue) in (D). The scale bars shown are 10µm.

We hypothesized that the R299Q variant–induced re-localization of the AMPKγ2 subunit during cardiogenesis may have downstream effects on mitochondrial and sarcomeric organization in cardiomyocytes. We used immunostaining with the Mitofusin-2 antibody to visualize mitochondria in adult ventricular sections. In TuAB and Tg^WT^ ventricles, mitochondria displayed a distinct alignment, forming longitudinal rows between myofibrils (**Supplement Figure 6A**). However, in Tg^R299Q^ ventricular myocytes, mitochondria appeared clumped and exhibited a less organized structure (**Supplement Figure 6A**). High-resolution TEM images further supported these observations. TuAB and Tg^WT^ ventricular sections exhibited a dense alignment of mitochondria alongside myofilaments, whereas Tg^R299Q^ ventricular sections showed clusters of mitochondria that were visibly separated from the myofibrils (**Supplement Figure 6B**). Furthermore, mitochondria were significantly enlarged in Tg^R299Q^ ventricular sections compared with TuAB and Tg^WT^ (**Supplement Figure 6C-D**). These results suggest that the R299Q variant disrupts both sarcomeric and mitochondrial organization and alters mitochondrial morphology in mature cardiomyocytes.

### The R299Q variant enhanced the physical interaction between AMPKγ2 and myosin

As the R299Q variant changes the localization of AMPKγ2 to myofibrils and alters the contractile function of myocytes, we hypothesized that there may be potential direct interactions between the AMPKγ2 and sarcomere proteins. We tested physical interactions between AMPKγ2 and a major sarcomere protein, myosin, with in situ proximity ligation assay (PLA) and co-immunoprecipitation methods. In adult ventricular sections, AMPKγ2 and myosin heavy chain antibodies generated positive PLA signals in both TuAB and Tg^WT^ hearts, indicating that AMPKγ2 and myosin reside in close proximity and potentially form a complex. Notably, Tg^R299Q^ ventricles exhibited substantially increased PLA signals compared to TuAB and TgWT ventricles, with approximately 5-fold and 2-fold increases, respectively (**Figure 6A, C**), suggesting that the R299Q variant strengthens the interaction between AMPKγ2 and myosin. We also performed immunoprecipitation from adult heart lysates using AMPKγ2 antibody-conjugated beads, followed by western blot analysis of the eluate fractions. Consistent with the PLA results, co-immunoprecipitation demonstrated an interaction between AMPKγ2 and myosin in both TuAB and Tg^WT^ hearts, which was greatly enhanced in Tg^R299Q^ hearts (**Supplement Figure 7**).

**Figure 6.**
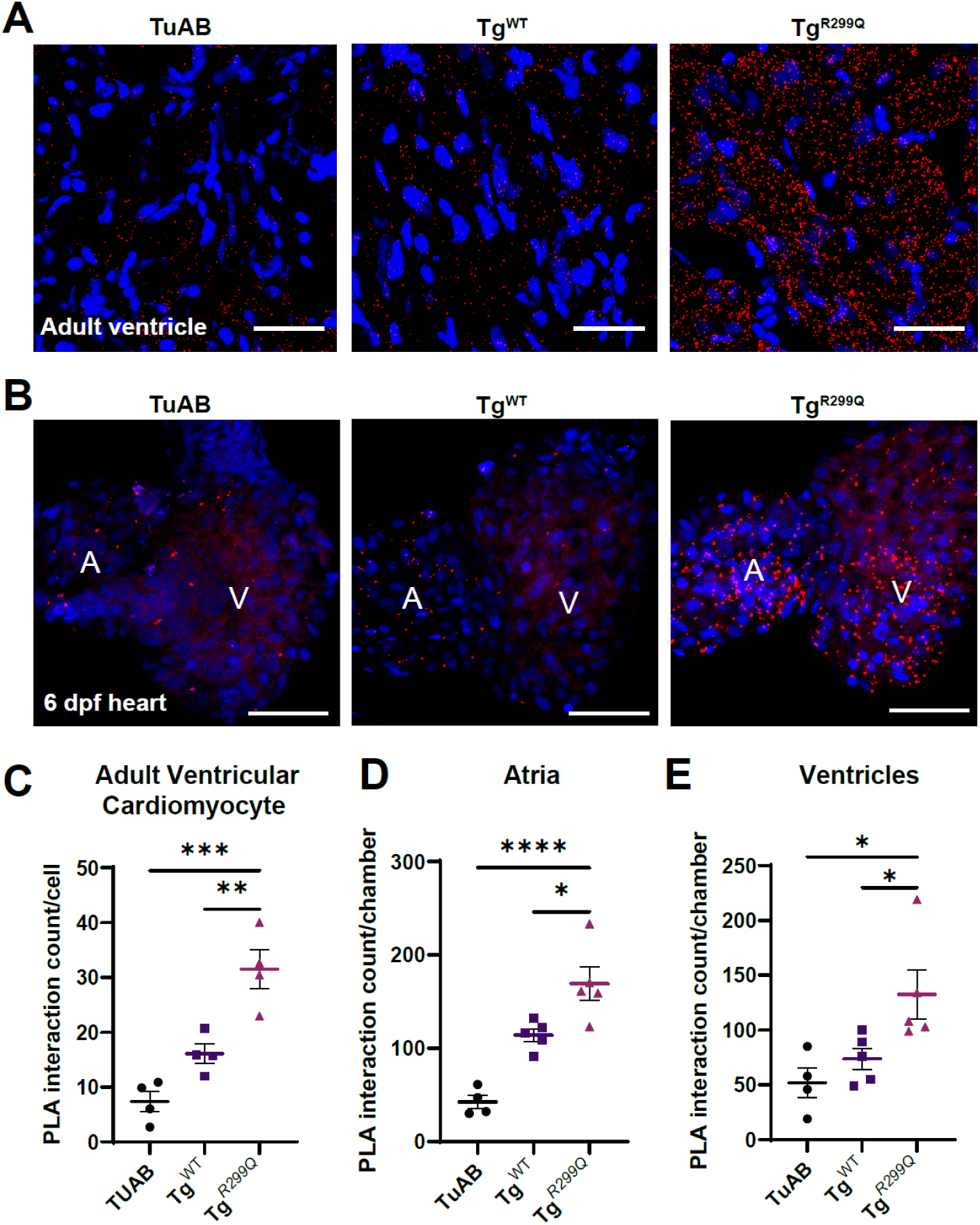
AMPKγ2 forms a macromolecular complex with myosin, and the R299Q variant enhances this interaction. A-B. Proximity ligation assays (PLAs) using antibodies for AMPKγ2 (rabbit) and myosin heavy chains (mouse). Shown are representative images from ventricular sections of TuAB, Tg^WT^, and Tg^R299Q^ adult hearts (A), and whole hearts isolated from 6 dpf TuAB, Tg^WT^, and Tg^R299Q^ larvae (B). The scale bars shown are 25µm. C. Quantified PLA interactions in ventricular sections of TuAB, Tg^WT^, and Tg^R299Q^ adult hearts. D-E. Quantified PLA interaction in atria (E) and ventricles (F) of TuAB, Tg^WT^, and Tg^R299Q^ 6 dpf hearts. Each data point in C-E represents an individual heart studied. *P < 0.05, **P < 0.01, ***P < 0.001, and ****P < 0.0001.

We next examined whether this interaction occurs during early cardiac development. In 6 dpf hearts, PLA detected AMPKγ2-myosin interactions in all three genotypes (**Figure 6B, D, E**). Similar to adult hearts, Tg^R299Q^ 6dpf hearts showed increased numbers of PLA puncta in both the atria and ventricles compared to TuAB and Tg^WT^ (**Figure 6B, D, E**). In contrast, the R299Q variant did not alter the interaction between AMPKγ2 and AMPK α subunits (**Supplementary Figure 8**).

### Altered Na⁺/Ca²⁺ exchanger (NCX) dependence of Ca^2+^ handling in Tg^R299Q^ hearts

To further define the mechanism underlying prolonged Ca^2+^ transient duration in Tg^R299Q^ hearts, we examined Ca^2+^ extrusion pathways. Because Tg^R299Q^ hearts exhibited prolonged Ca^2+^ decay without changes in peak transient amplitude or diastolic Ca^2+^, this phenotype is more consistent with altered Ca²⁺ removal kinetics than with increased cellular Ca^2+^ load or enhanced SR Ca^2+^ release. To test whether NCX-dependent extrusion contributes to this defect, we assessed the effects of NCX inhibitor SEA0400 on Ca^2+^ dynamics in Tg^WT^ and Tg^R299Q^ hearts. In Tg^WT^ hearts, 50 nM SEA0400 prolonged Ca^2+^ duration (at 80% decay) (**Figure 7A-B**) and increased diastolic Ca^2+^ (**Figure 7C-D**) in the ventricles, consistent with a contribution of forward-mode NCX to Ca²⁺ extrusion. In contrast, SEA0400 did not alter Ca²⁺ duration or diastolic Ca^2+^ in Tg^R299Q^ hearts (**Figure 7A-D**), suggesting that NCX-dependent extrusion makes a limited contribution to Ca^2+^ decay in the mutant. RNA-seq showed no change in NCX transcript expression (adjusted P-value=0.90). Notably, SEA0400 increased Ca^2+^ transient amplitude in both atria and ventricles of Tg^R299Q^ hearts, whereas in Tg^WT^ hearts this effect was restricted to the atria (**Figure 7E-F**). These findings indicate that NCX remains functionally engaged in Tg^R299Q^ hearts, but its contribution to Ca^2+^ decay is diminished, suggesting that another process, potentially increased myofilament Ca^2+^ buffering, becomes rate-limiting for Ca^2+^ removal in Tg^R299Q^ (**Figure 7G**).

**Figure 7.**
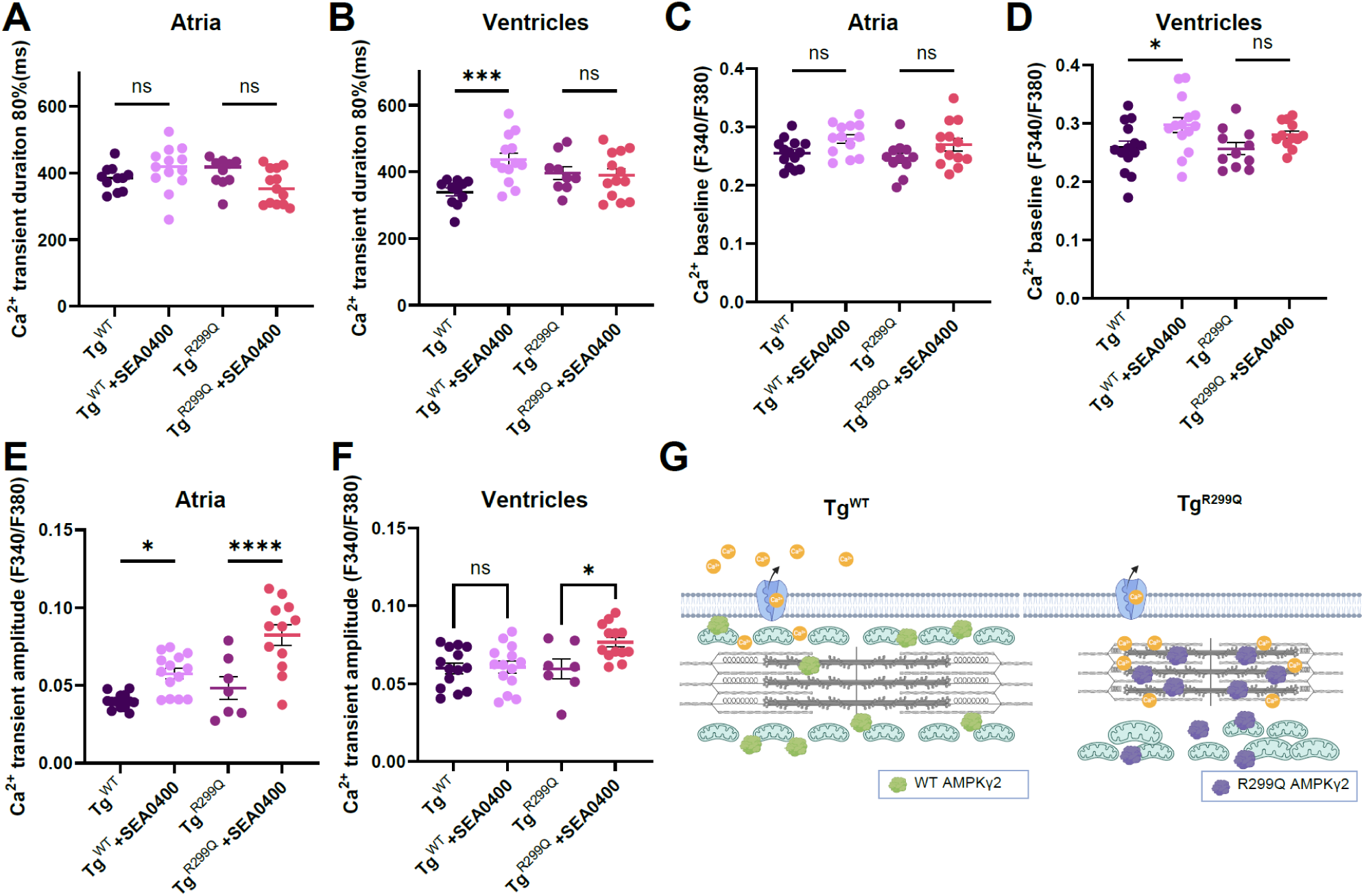
NCX inhibition differentially affects Ca^2+^ dynamics in Tg^WT^ and Tg^R299Q^ hearts. A-B. Duration of the Ca^2+^ transient (at 80% decay) in atria (A) and ventricles (B) of 6 dpf Tg^WT^ and Tg^R299Q^ hearts under DMSO control or treated with NCX blocker SEA0400 (50 nM, 1 hour). [Ca^2+^]_i_ was calculated by fluorescence ratio of Fura2 (F340/F380). C-D. Diastolic Ca^2+^ (baseline) in atria (C) and ventricles (D) of 6 dpf Tg^WT^ and Tg^R299Q^ hearts under DMSO control or treated with SEA0400. E-F. The amplitude of Ca^2+^ transient, which is measured as the difference between systolic and diastolic [Ca^2+^]_i_ in atria and ventricles of 6 dpf hearts, comparing Tg^WT^ and Tg^R299Q^ hearts under DMSO control or treated with SEA0400. G. Schematic illustrating a potential mechanism by which enhanced AMPKγ2-myosin interaction by R299Q increases myofilament Ca^2+^ retention, thereby limiting NCX-mediated Ca^2+^ extrusion. Results are expressed as mean ± S.E.M. Each data point represents the mean of an individual heart studied. *P < 0.05, ***P < 0.001, and ****P < 0.0001.

### Inhibiting myosin decreased AMPKγ2-myosin interactions and rescued electrical disturbances caused by R299Q

We reasoned that, if enhanced AMPKγ2-myosin interaction and myofilament Ca^2+^ retention are causal for the electrophysiological abnormalities in Tg^R299Q^, then modulating myosin should ameliorate these phenotypes. To test this, we used both pharmacological and genetic approaches.

We treated 2 dpf larvae with 2 µM mavacamten, a myosin ATPase inhibitor, and assessed its effects at 6 dpf. Notably, mavacamten reduced the number of PLA puncta between AMPKγ2 and myosin (**Figure 8A-C**), suggesting that myosin activity or conformation modulates AMPKγ2-myosin interaction. In Tg^R299Q^ hearts, reduced AMPKγ2-myosin interaction after mavacamten treatment was associated with increased ventricular conduction velocity (**Figure 8D, J**) and shortening of the prolonged atrial APD (**Figure 8K**), although ventricular APD was not significantly shortened (**Figure 8L**). In contrast, mavacamten did not significantly affect these electrophysiological parameters in Tg^WT^ hearts, indicating a preferential effect on the R299Q hearts(**Figure 8E-H**).

**Figure 8.**
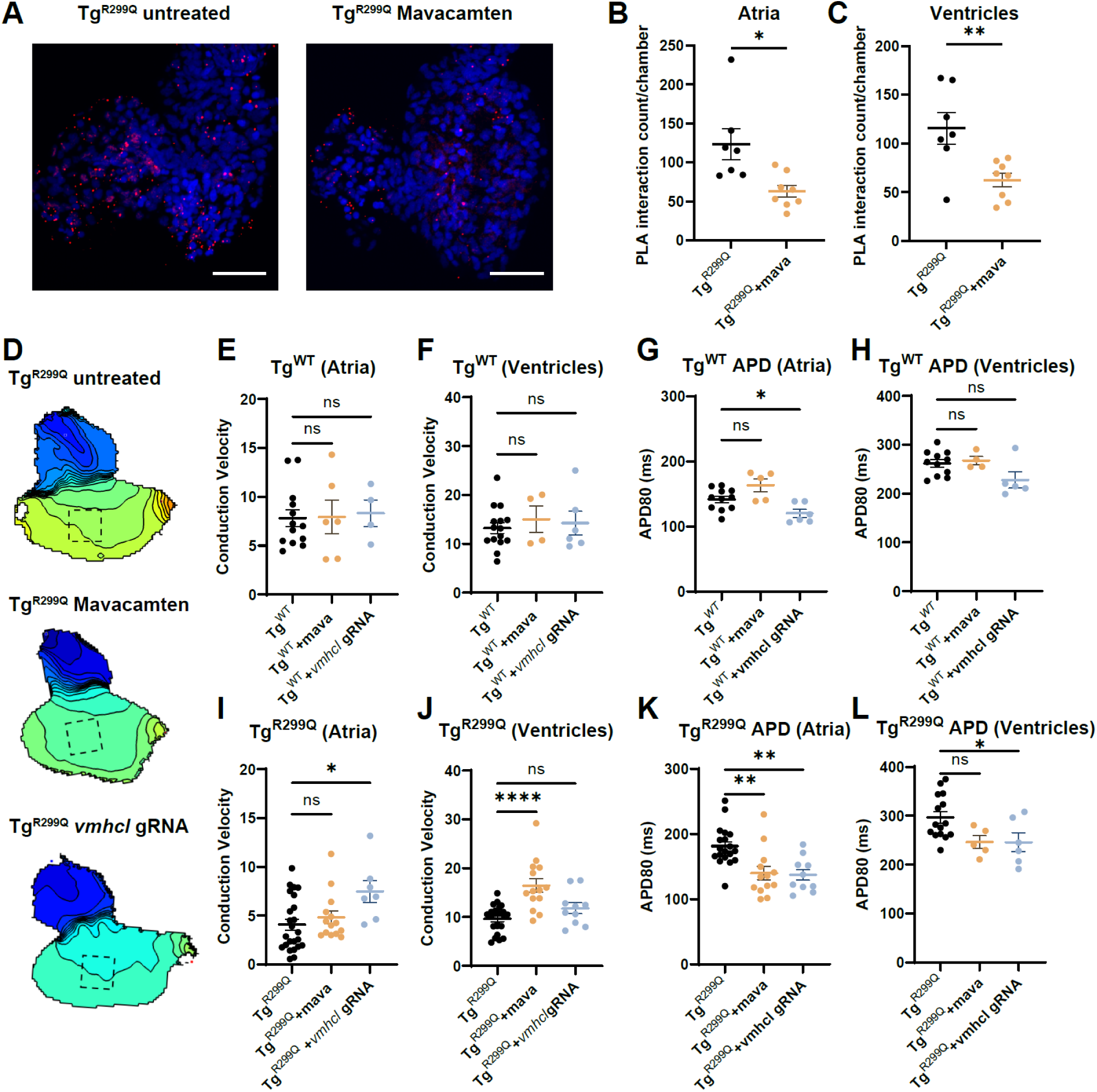
Inhibition of cardiac myosin decreases AMPKγ2-myosin interaction and restores early embryonic electrophysiological abnormalities induced by the R299Q variant. A. PLAs using antibodies for AMPKγ2 (rabbit) and myosin heavy chains (mouse). Shown are representative images from whole hearts isolated from 6 dpf Tg^R299Q^ larvae, under control and 2µM mavacamten conditions. The scale bars shown are 25µm. B-C. Quantified PLA interaction in atria (B) and ventricles (C) of Tg^R299Q^ 6 dpf hearts under control and mavacamten conditions. D. Example isochrone maps of hearts isolated from 6 dpf Tg^R299Q^ fish, comparing the untreated, treatment of myosin inhibitor Mavacamten, and *vmhcl* gRNA knockdown conditions. E, F, I, J. Conduction velocity of the atria and ventricles from 6 dpf hearts of Tg^WT^, Tg^WT^ treated with Mavacamten, Tg^WT^ injected with *vmhcl*-targeting gRNA, Tg^R299Q^, Tg^R299Q^ treated with Mavacamten, and Tg^R299Q^ injected with *vmhcl*-targeting gRNA. The hearts were labeled with FluoVolt and analyzed using optical mapping. G, H, K, L. Measured AP duration of the atria and ventricles from 6 dpf hearts of Tg^WT^, Tg^WT^ treated with Mavacamten, Tg^WT^ injected with *vmhcl*-targeting gRNA, Tg^R299Q^, Tg^R299Q^ treated with Mavacamten, and Tg^R299Q^ injected with *vmhcl*-targeting gRNA. The hearts were paced at 80 beats per minute for the quantification of the APD_80_. Results are expressed as mean ± S.E.M. Each data point represents the mean of an individual heart studied. *P < 0.05, **P < 0.01, and ****P < 0.0001.

In parallel, we used CRISPR/Cas9 to knock down *vmhcl*, the zebrafish ortholog of human *MYH7*^19^. The designed gRNA targets Exon 2 of *vmhcl*, generating mosaic loss of function edits. We found that the knockdown of *vmhcl* significantly increased the conduction velocity in the atria of the Tg^R299Q^ 6 dpf larvae (**Figure 8D, I**), restoring the conduction velocity to levels comparable to the Tg^WT^ (**Figure 8E**). No changes in the ventricular conduction velocity were observed in the *vmhcl* knockdown (**Figure 8F, J**). Additionally, *vmhcl* knockdown reduced the level of APD prolongation caused by R299Q in both atria and ventricles (**Figure 8K, L**). Together, these results indicate that myosin inhibition reduces AMPKγ2-myosin interaction and partially rescues the early electrophysiological abnormalities in Tg^R299Q^ larval hearts.

## Discussion

*PRKAG2* cardiac syndrome is also known as a glycogen storage disease since intracellular glycogen accumulation is the most noticeable cellular phenotype observed in the hearts of affected individuals. However, it is important to recognize that this glycogen buildup may be a downstream consequence rather than the primary disease mechanism. Therefore, it is crucial to identify the molecular mechanisms underlying *PRKAG2* variants to effectively manage their associated cardiomyopathy and conduction disorders. In this study, we investigated the impact of a *PRKAG2* variant on cardiomyocyte cytoarchitecture, morphology, and electrophysiology, through the process of cardiogenesis, with a focus on its role in the pathogenesis of HCM. Our examination of the heart during early development unveiled remarkable electrical disturbances and alterations in AMPKγ2 subcellular localizations induced by the R299Q variant, all of which occurred prior to the onset of any detectable glycogen accumulation. This finding suggests that the initial pathogenesis of *PRKAG2* cardiomyopathy takes place during cardiogenesis, independent of the well-established glycogen storage effect. We further identified a physical interaction between AMPKγ2 and myosin that was enhanced by the R299Q variant. This altered AMPKγ2-myosin interaction potentially affects myosin function and myofilament Ca²⁺ buffering, thereby contributing to the observed electrophysiological abnormalities. Our findings provide new mechanistic insight into the pathogenesis of *PRKAG2* syndrome.

### The dynamic effect of *PRKAG2* variants on AMPK phosphorylation

Previous investigations into *PRKAG2* pathological variants have demonstrated variable effects on AMPK phosphorylation^6,12,20–22^. Indeed, even the specific variant in the current study, R302Q (R299Q in the murine isoform), has been shown to exhibit either up– or down-regulation of AMPK phosphorylation across different experimental systems^7,21^. Moreover, the timing of expression has been found to influence these effects, with chronic overexpression of R302Q resulting in decreased AMPK phosphorylation, while transient overexpression leads to increased AMPK phosphorylation^23^. Our results revealed a substantial reduction in AMPK phosphorylation caused by the R299Q variant during early cardiogenesis. This effect became less pronounced in adult hearts, suggesting that the *PRKAG2* variant dynamically modulates AMPK activity throughout cardiac development. *PRKAG2* variants were thought to prevent AMPK from redirecting metabolism towards increased catabolism and decreased anabolism, resulting in glycogen accumulation. However, previous studies and our own results suggest that *PRKAG2* variants elevate glycogen storage in mature cardiomyocytes, regardless of the directionality of AMPK modulation^12,23^. These results support the notion that glycogen accumulation resulting from *PRKAG2* mutation is a gradual process occurring over time, stemming from the dysregulation of AMPK responses throughout cardiac development and beyond. In addition to its metabolic roles, AMPK activation can restrain protein synthesis via mTORC1 inhibition and promote autophagy through ULK1 activation^24^. Dysregulation of these processes by the R299Q variant may therefore represent additional mechanisms contributing to cardiac hypertrophy beyond those described here.

### *PRKAG2* modulates cellular electrophysiology and conduction system development

Patients diagnosed with PRKAG2 cardiomyopathy commonly exhibit cardiac arrhythmias, such as ventricular pre-excitation, atrial fibrillation, conduction deterioration, and ventricular arrhythmias^8^. We demonstrated several cardiac electrophysiological changes during early embryogenesis in our *Prkag2* transgenic models. The slowed conduction velocity, together with prolonged APD and Ca^2+^ transient duration in atria and ventricles, can contribute to the development of atrial fibrillation and ventricular arrhythmias. Our data suggest that enhanced interaction of the mutant γ2 subunit with myosin may alter Ca^2+^ binding and dissociation from the myofilament, thereby prolonging Ca^2+^ transient duration and APD. Inhibition of myosin with mavacamten reduced mutant AMPKγ2-myosin interaction and alleviated APD prolongation and conduction abnormalities in Tg^R299Q^ hearts, supporting a causal role for altered AMPKγ2-myosin interaction in the electrophysiological phenotype. Changes in ion channel expression (**Supplementary Figure 4**) may also contribute to these electrical abnormalities. Further studies are needed to define the mechanisms underlying slowed conduction and to determine whether related mechanisms also contribute to the ventricular pre-excitation phenotype observed in *PRKAG2* syndrome.

### AMPKγ2 balances energy production and consumption in cardiomyocytes

The AMPKγ2 subunit functions as a cellular energy sensor through adenine nucleotide binding and regulation of AMPK activity. Our findings show that AMPKγ2, particularly the mutant form, interacts with myosin and localizes near the myofilament, suggesting that it may serve as a local energy sensor within the sarcomere. In addition to sensing ATP availability, AMPKγ2 may influence contractile function through its interaction with myosin, thereby affecting the rate of ATP consumption in cardiomyocytes. In the Tg^R299Q^ model, we observed increased sarcomere contractility, which is likely to elevate ATP demand and increase ADP/ATP and AMP/ATP ratios. At the same time, the R299Q variant impairs AMPK activation partly because of reduced nucleotide-binding affinity relative to the wild type^25^, limiting activation of the catabolic pathways needed to restore energy balance. Thus, increased ATP consumption coupled with impaired metabolic compensation may create a sustained energetic deficit in cardiomyocytes. Over time, this imbalance could contribute to progressive cardiac dysfunction and eventual heart failure, as observed in aged mutant fish and in patients with *PRKAG2* syndrome.

### Implications of the direct interaction between the AMPKγ2 and myosin

It is well established that sarcomere contraction and cellular metabolism are closely intertwined^26^. In HCM caused by sarcomere gene mutations, inefficient contraction leads to increased ATP consumption and ADP elevation, ultimately resulting in mitochondrial dysfunction^9,26^.

Our study demonstrated a direct interaction between an energetic sensor and a myofilament protein. Although the molecular basis of this interaction remains to be defined, we observed a stronger interaction induced by the *Prkag2* R299Q variant. This enhanced interaction has the potential to influence myosin’s localization, functions, and turnover. In addition, we found that the myosin inhibitor mavacamten reduces the AMPKγ2-myosin interaction in the R299Q mutant, suggesting that this interaction is dynamic and may depend on myosin conformational or functional state. Likewise, it is plausible that HCM-causing variants in *MYH7* could reciprocally affect AMPKγ2 functions, thereby altering AMPK activity and shifting cellular metabolism. Notably, prior studies have shown that AMPKα phosphorylates cardiac troponin I, increasing contractility and Ca²⁺ sensitivity^27,28^. Further studies are needed to determine whether the AMPKγ2-myosin interaction recruits the AMPK holoenzyme to the myofilament, enabling local catalytic effects on targets such as troponin I. Our findings suggest a distinct mechanism underlying intricate crosstalk between sarcomere contraction and metabolism that extends beyond energy demand. Moreover, they also unveil a partially overlapping mechanism underlying sarcomere-linked and *PRKAG2*-associated HCMs.

### Limitations

In this study, we utilized transgenic models expressing murine *Prkag2* instead of directly editing the zebrafish orthologs of *PRKAG2*, since zebrafish possess multiple orthologs of *PRKAG2* (*prkag2a* and *prkag2b*), which require CRISPR-Cas9 editing in both genes. Our approach has limitations, including potential species-specific and overexpression effects. To account for transgenesis-related effects, we generated a WT *Prkag2* transgenic line as a control for the R299Q model. Although human, murine, and zebrafish PRKAG2 isoforms are highly conserved, particularly in the CBS domains, future studies in complementary model systems will be needed to further validate our findings.

Additionally, several differences between zebrafish and human cardiac electrophysiology should be considered when interpreting our findings. Unlike the human ventricle, which relies predominantly on NaV1.5-mediated Na^+^ current for depolarization, zebrafish ventricular action potentials also depend on T-type Ca^2+^ currents^29^. This makes the zebrafish heart inherently more sensitive to myofilament Ca^2+^ retention. Moreover, zebrafish lack a discrete His-Purkinje system, with ventricular conduction appearing to rely on gap junction-mediated cell-to-cell coupling through trabecular cardiomyocytes^30^. Despite these differences, the key molecular processes underlying the mechanisms we describe are highly conserved between zebrafish and humans.

In summary, our study uncovered novel mechanisms driving the early pathogenesis of *PRKAG2*-linked cardiomyopathy that are distinct from the well-established glycogen storage effect. These mechanisms revealed how a cellular energy sensor can influence the electrical and contractile functions of myocytes. Our findings open up new therapeutic avenues for addressing *PRKAG2*-linked HCM and potentially offer insights into treating other prevalent forms of cardiomyopathies.

## Acknowledgements

This work was supported by the National Institutes of Health (5R01HL164675 and 5R24OD035402) and American Heart Association (24IPA1268047). We thank Dr. Olivia Weeks and the Harvard Medical School Electron Microscopy Facility for assistance with experiments.

## Materials and Methods

### Animal models

The Tg (*cmlc2*:*Prkag*2^WT^) and Tg (*cmlc2*:*Prkag*2^R299Q^) fish lines were generated with a Tol2 transposase system. The *cmlc2* promoter, murine *Prkag2* cDNA with or without the R299Q mutation, and SV40 poly (A) sequences were inserted into the destination vector pDestTol2pACryGFP-containing Tol2 sequence, and the transgene *cryaa*:GFP that drives enhanced GFP expression in the eyes. The plasmid was then co-injected with transposase mRNA into zebrafish embryos at the one-cell stage. Founders (F0) with GFP+ eyes were then outcrossed with the TuAB fish to establish a stable transgenic F1 line. Copy numbers of the transgenes were quantified using the comparative Ct method ^31^.Genomic DNA was extracted from fin clips of adult fish and subjected to qPCR using a set of primers for GFP and the control gene ^31^. Δ(Ct) was used to calculate the copy number. Animals with 2-3 copies of the transgene were used for experiments. All adult animals used for the sample set of experiments were controlled for age and sex.

The *vmhcl* knockdown zebrafish was generated with the acute CRISPR-Cas9 method. The guide RNA was designed to target “AGCCACCGTCGTGAGCCGAG” in Exon 2 of *vmhcl*. The single guide RNA (sgRNA) was synthesized (Integrated DNA Technology) and incubated with Alt-R Cas9 Nuclease from the ribonucleoprotein (RNP) complex. 1.5 nl of the RNP complex was then injected into zebrafish embryos at the one-cell stage. Sanger sequencing was performed on zebrafish larvae used for experiments to ensure >30% editing at the targeted site.

### Echocardiography

Adult zebrafish were anesthetized with 0.02% tricaine (MS-222) until no movement, then transferred to the imaging chamber containing system water with 0.02% tricaine. The fish were positioned ventral side up using a pre-shaped sponge. Echocardiography was performed using the Vevo3100 ultrasound imaging platform (Visual Sonics/Fujifilm), equipped with a high-frequency transducer (MX700). The transducer was placed longitudinally and proximal to the ventral side of the fish. The relative positions of the fish and transducer head were adjusted using a micro-manipulator. Two-dimensional (B-mode) images containing >6 cardiac cycles were recorded. Image acquisition of each fish was completed within 2 minutes. After recording, fish were transferred to the recovery tank, where they generally recovered within 30 seconds with no death observed. Echo images were analyzed using ImageJ. The ejection fraction (EF) was calculated based on end-diastolic volume (EDV) and end-systolic volume (ESV) according to EF=(EDV–ESV)/EDV; EDV=(8*Diastole_Area^2^)/3*π*Diastole_long_axes; ESV=(8*Systole_Area^2-^)/3*π*Systole_long_axes. The EDVs were not corrected by body size/weight, as the fish of each genotype were comparable in size. Both female and male zebrafish were used for the echo analyses.

### Quantification of heart contractile function in zebrafish larvae

The ventricular contractile function of zebrafish larvae was quantified as previously described^32^. Briefly, 3 dpf fish larvae were transferred to the imaging chamber and allowed to acclimate to microscope illumination for 1 minute. Videos of the ventricle from the lateral view were recorded using the Nikon upright microscope and the FASTCAM Mini AX100 camera (Photron). Images were acquired at 250 frames per second (fps) for 3 seconds, encompassing ∼6 cardiac cycles with a Plan Apo 20x/0.75 objective. Acquired images were subsequently analyzed using ImageJ. The ventricular volume was calculated based on v=4/3*π*long_axes*short_axes^2^. The cardiac output was calculated as stroke volume*beats per minute.

### Glycogen Assay and Histochemistry

The glycogen content in adult and embryonic hearts was quantified using the Glycogen Fluorometric Assay Kit (Cayman Chemical) according to the manufacturer’s directions. Briefly, hearts from adult zebrafish were isolated, flash-frozen, weighed, and homogenized in the assay buffer containing a protease inhibitor cocktail. 50µl assay buffer was used for each 1 mg of heart tissue. The fluorescence of the assay plate was measured at an excitation wavelength of 530-540 nm and an emission wavelength of 585-595 nm using Cytation 5 (Biotek). For 6 dpf embryonic hearts, 10 hearts were isolated and pooled into one sample, and 50µl assay buffer was used for every 10 hearts. The glycogen content of the samples was calculated based on the standard curve.

Periodic Acid-Schiff (PAS) staining was performed using the PAS staining system (Sigma Aldrich) according to the manufacturer’s instructions. Isolated 6 dpf hearts were fixed with 4% PFA prior to PAS staining. Samples were not counterstained with Hematoxylin, as the nuclear size to cell size ratio in the heart is very high at this developmental stage, and staining with Hematoxylin would limit the visualization of PAS stain intensity.

### Immunofluorescence

Embryonic hearts were isolated from 3 or 6 dpf larvae in Tyrode’s solution and fixed with Prefer fixative (Anatech) for 15 minutes at room temperature. They were then washed, blocked, and incubated with primary antibodies at 4 °C overnight. The secondary antibodies are Alexa 488-conjugated goat anti-rabbit antibody and Alexa 546-conjugated goat anti-mouse antibody. Confocal images were acquired using an Olympus FV1200 Confocal Microscope and a Nikon AXR Live-Cell Point-Scanning Confocal Microscope and analyzed using ImageJ.

### Proximity ligation assay (PLA)

The in situ proximity ligation assay was performed using the Duolink in Situ Orange fluorescence kit (Sigma) in accordance with the manufacturer’s instructions. Briefly, the 4% PLA fixed heart samples were initially blocked using Duolink blocking reagent and subsequently incubated with primary antibodies at 4°C overnight. After washing, the samples were then exposed to PLA probes at 37°C for 1 hour, followed by ligation and amplification steps. To serve as negative controls, samples were solely incubated with the AMPKγ2 antibody during the primary antibody incubation step. Confocal imaging settings were adjusted to ensure that the negative control samples did not display any PLA signals.

### Optical mapping

Hearts were isolated from 6 dpf embryos and placed in normal Tyrode’s solution (136 mM NaCl, 5.4 mM KCl, 1.0 mM MgCl_2_, 0.3 mM NaH_2_PO_4_, 1.8 mM CaCl_2_, 5 mM glucose and 10 mM HEPES, pH 7.4) with 1% bovine serum albumin. For voltage mapping, isolated hearts were incubated with FluoVolt membrane potential dye at 1:500 dilution in Tyrode’s solution for 15 minutes. The individual heart was then placed in a recording chamber containing Tyrode’s solution with 10 µM of Cytochalasin D to inhibit contraction. The excitation light was generated by Cairn OptoLED, transmitted through a 475/35-nm excitation filter, and reflected by a 506 nm cut-on dichroic mirror. The fluorescent images of 80 by 80-pixel resolution were acquired at a frame rate of 2000 frames/s as previously described^33^. For ratiometric Ca^2+^ mapping, isolated hearts were incubated in Tyrode’s solution containing 50 μM calcium-sensitive dye Fura-2 AM for 20 minutes, followed by washing and incubation in dye-free Tyrode’s solution for 45 minutes. Ratiometric Ca^2+^mapping was conducted using the imaging setup described previously. Images were acquired at a frame rate of 500 frames/s using the SciMeasure DaVinci 2K CMOS Camera. The hearts were paced with field stimulation at the rate of 1.33 Hz for Ca^2+^ mapping and action potential duration (APD) measurements.

Acquired images were analyzed with a customized MATLAB code as described previously^33^. The regions of interest (ROIs) for atrial conduction velocity and Vmax is 15 by 15 pixels, for ventricular conduction velocity and Vmax is 20 by 20 pixels, and for APD measurements is 10 by 10 pixels.

### Co-Immunoprecipitation

Protein samples for the co-immunoprecipitation (co-IP) assay were prepared by homogenizing excised adult zebrafish hearts on ice in co-IP buffer (50 mM HEPES (pH 7.4), 250 mM NaCl, 2 mM EDTA, 0.5% (v/v) NP-40 substitute, 10% Glycerol supplemented with cOmplete protease and phosphatase inhibitor). Lysates were centrifuged at 16200 rcf at 4°C for 15 mins. The supernatants were transferred to a fresh tube followed by total protein quantification with Pierce^TM^ BCA protein assay. 500 μg of lysates was incubated with 1 μg of anti-Rabbit IgG control or anti-PRKAG2 antibody (12568-1-AP; Proteintech) overnight at 4°C. Immune complexes were incubated with protein A/G plus agarose beads (sc-2003; Santa Cruz Biotechnology) for 3 hours at 4°C. The beads were washed three times in co-IP buffer and eluted in 2X sample loading buffer at 100 °C for 5 mins. The eluted samples and 10% of the input were separated by SDS-polyacrylamide gel electrophoresis. The separated proteins were transferred to 0.2 µm PVDF membrane (1620174; Biorad). The membranes were blocked with 5% non-fat milk (9999S; Cell Signaling) in Tris-buffered saline containing 0.1% Tween-20 (TBST) at room temperature for 1 hour. The membranes were incubated with the respective primary antibodies (dilution 1:1000) overnight at 4°C [anti-GAPDH (2118S; Cell Signaling), anti-myosin heavy chain (F59; DHSB)]. The next day, the membranes were washed three times followed by incubation with the respective peroxidase conjugated secondary antibodies (dilution 1:1000) for 1 hour at room temperature. All blots were developed using ProSignal Femto ECL reagent (20-302; Genesee scientific).

### Western blot

Protein samples for western blot analyses were prepared by homogenizing excised adult zebrafish hearts on ice in Pierce^TM^ RIPA buffer supplemented with cOmplete protease and phosphatase inhibitor. Lysates were centrifuged at 16200 rcf at 4°C for 15 mins. Total proteins were quantified as described above. 40 µg of lysates were loaded onto the SDS-polyacrylamide gel. The protein transfer, membrane blocking, antibody incubation, and detection were performed as described above.

### Transmission Electron Microscopy

Freshly dissected adult zebrafish hearts were washed with Tyrode’s solution and then fixed in 5.0% glutaraldehyde, 2.5% paraformaldehyde, and 0.03% picric acid in 0.1 M sodium cacodylate buffer (pH 7.4) for 24 hours at 4 °C and embedded in resin. The samples were then sectioned on a Reichert Ultracut S Microtome to obtain ultrathin sections. The sections were then stained with lead citrate and imaged using a JEOL 1200 EX transmission electron microscope with an AMT 2k CCD camera. Representative images and images used for quantification of mitochondria and sarcomere morphology were taken close to the apex of the ventricle.

### RNA sequencing and analysis

Ventricles were isolated from dissected adult hearts and homogenized in TRIzol. RNA was extracted and purified using TRIzol RNA purification protocol. Ensembl release 101 transcriptome and genome data were used for alignment-free quantification of sequencing reads with Salmon v1.4.0^34^. Transcript-level counts were mapped to genes using tximeta, and DESeq2 was used for differential expression analysis^35^. Benjamini-Hochberg–adjusted P values of <0.05 were deemed as significant and used for gene set enrichment analyses, which were performed with clusterProfiler.

### Statistical Analysis

Blinding was implemented where possible throughout the study. Echocardiography, cardiac output, and PLA were performed and analyzed blinded to fish genotype. Optical mapping analyses were conducted using an automated analysis program, inherently removing bias. Samples were excluded from analysis if signal quality was poor or if they failed the quality control criteria predefined by the automated program.

All statistical analyses were performed with GraphPad Prism 9 or MATLAB. The normality of the data was first assessed with Kolmogorov-Smirnov test. If the distribution was normal, we then used unpaired Student’s t tests. If the distribution was not normal, we used the nonparametric Mann-Whitney test. For comparisons among more than two groups, we used ordinary one-way analysis of variance (ANOVA) tests followed by Dunnett’s multiple comparison adjustment. A P value or adjusted P value of <0.05 was used as the threshold to determine statistical significance.

## Figures and Figure Legends

**Supplement Figure 1.**
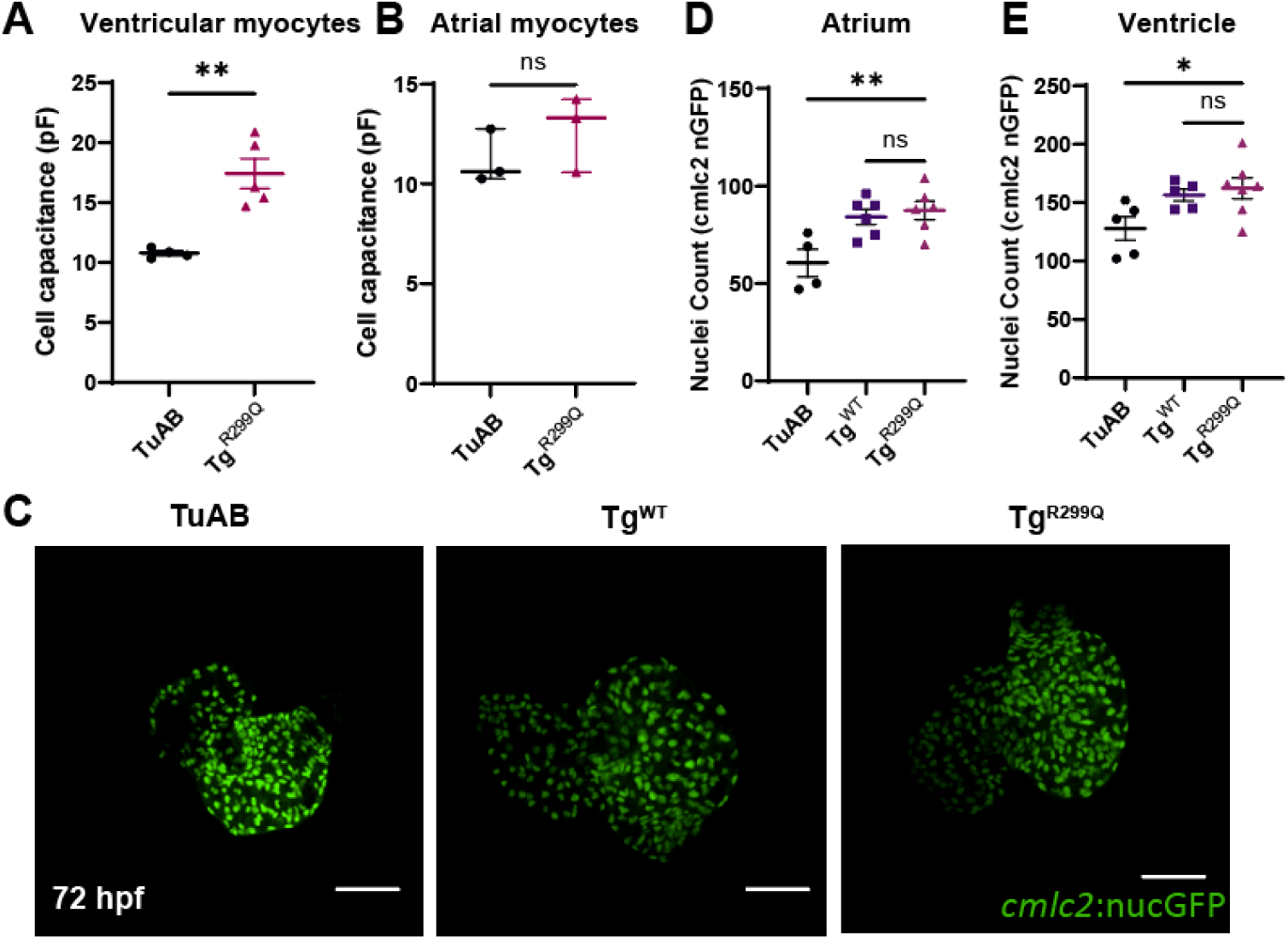
Cardiac transgenic expression of AMPK γ2 alters the cardiomyocyte number and membrane capacitance. A-B. Measurement of cell membrane capacitance using the patch clamp whole-cell configuration of isolated ventricular and atrial cardiomyocytes from TuAB and Tg^R299Q^ adult hearts. Each data point represents a cell, and two animals from each genotype were used for the analysis. C. Confocal Z-stack projections of isolated hearts from TuAB, Tg^WT^, and Tg^R299Q^ larvae, in the background of the myocardial nuclear reporter line Tg(*cmlc2*:nucGFP). GFP+ nuclei indicate nuclei of cardiomyocytes. D-E. Quantification of the cardiomyocyte nuclei number in atria (D) and ventricles (E) of 3 dpf TuAB, Tg^WT^, and Tg^R299Q^ hearts. Results are expressed as mean ± S.E.M. *P < 0.05 and **P < 0.01.

**Supplement Figure 2.**
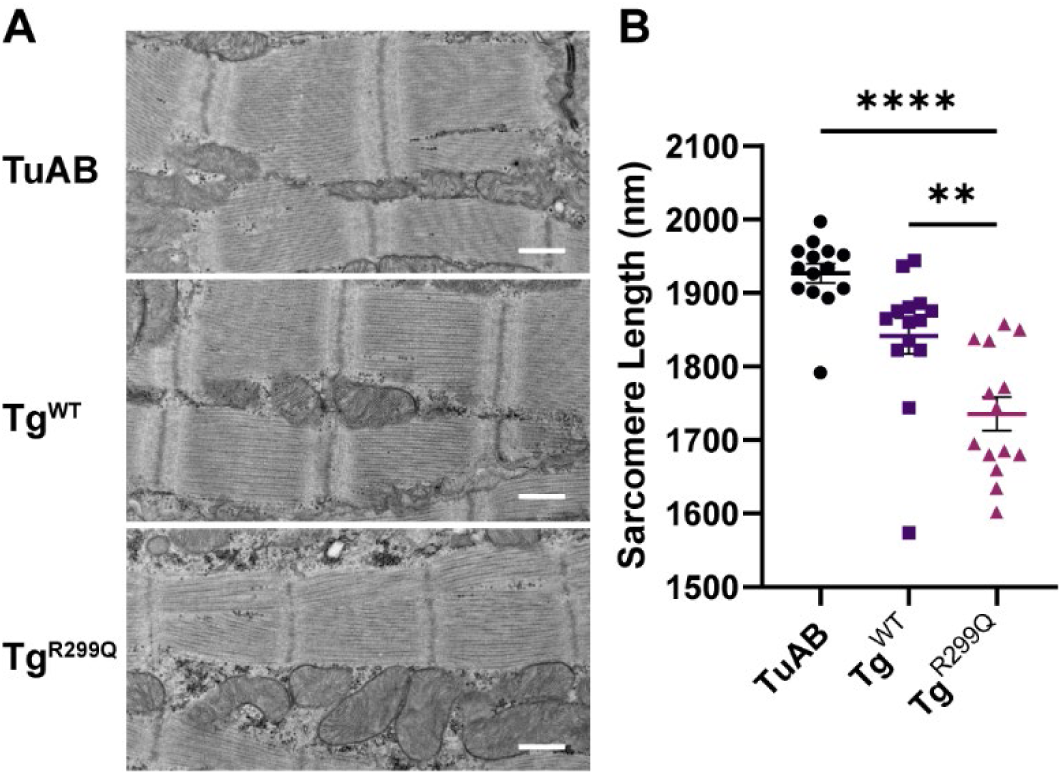
Tg^R299Q^ adult cardiomyocytes exhibited decreased sarcomere length. A. Representative TEM images of longitudinal ventricular sections of the TuAB, Tg^WT^, and Tg^R299Q^ adult hearts. Scale bars measure 500 nm. B. Quantified sarcomere length, measured as distances between adjacent z-disks, of ventricular sections of the TuAB, Tg^WT^, and Tg^R299Q^ adult hearts. Each data point represents the mean of 3 sarcomere units. Four heart samples of each genotype were used for this analysis.

**Supplement Figure 3.**
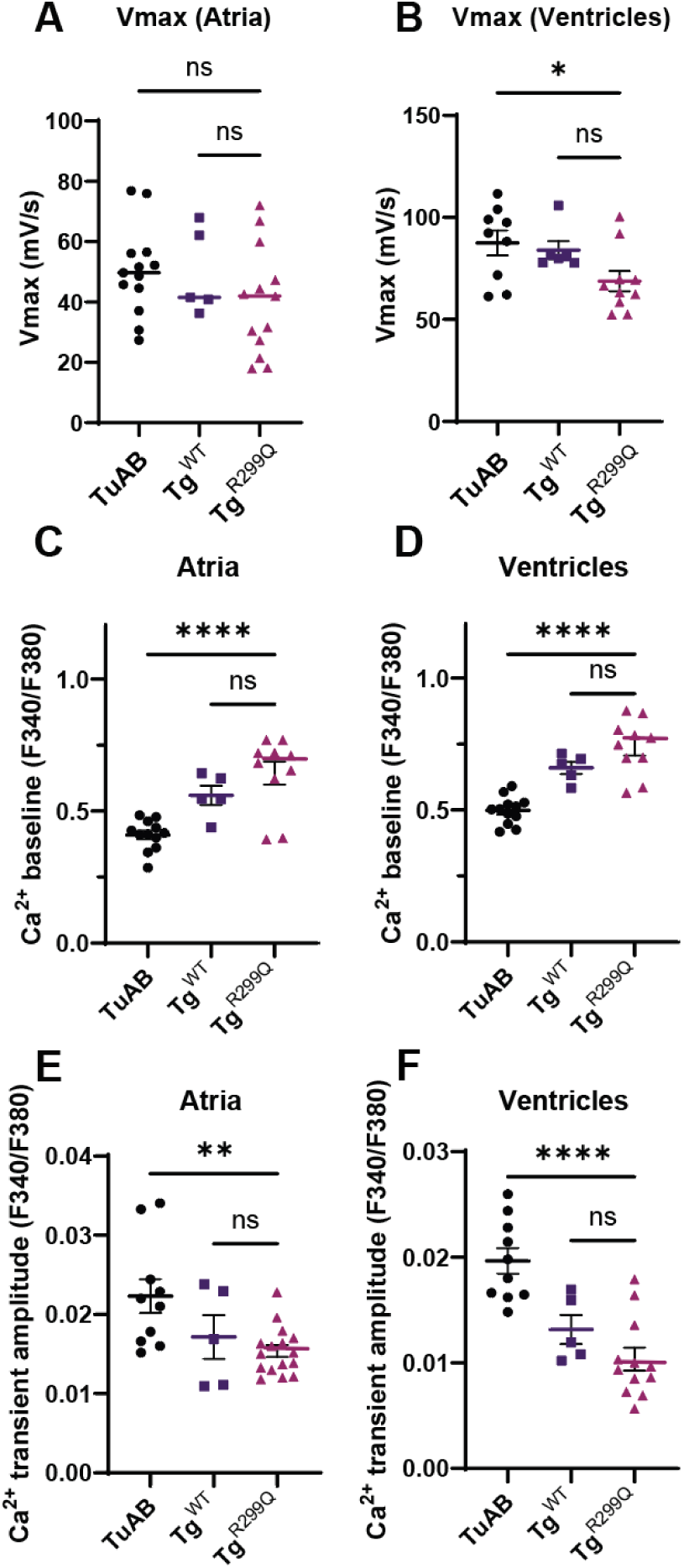
Other changes in the electrical behaviors of Tg^WT^ and Tg^R299Q^ hearts at 6 dpf. A-B. The maximum upstroke velocity of action potentials in atria and ventricles of 6 dpf hearts comparing TuAB, Tg^WT^, and Tg^R299Q^. C-D. The diastolic [Ca^2+^]_i_ (baseline) of atria and ventricles of 6 dpf hearts comparing TuAB, Tg^WT^, and Tg^R299Q^. E-F. The amplitude of Ca^2+^ transient, which is measured as the difference between systolic and diastolic [Ca^2+^]_i_ in atria and ventricles of 6 dpf hearts comparing TuAB, Tg^WT^, and Tg^R299Q^. Results are expressed as mean ± S.E.M. Each data point represents the mean of an individual heart studied. *P < 0.05, **P < 0.01, and ****P < 0.0001.

**Supplement Figure 4.**
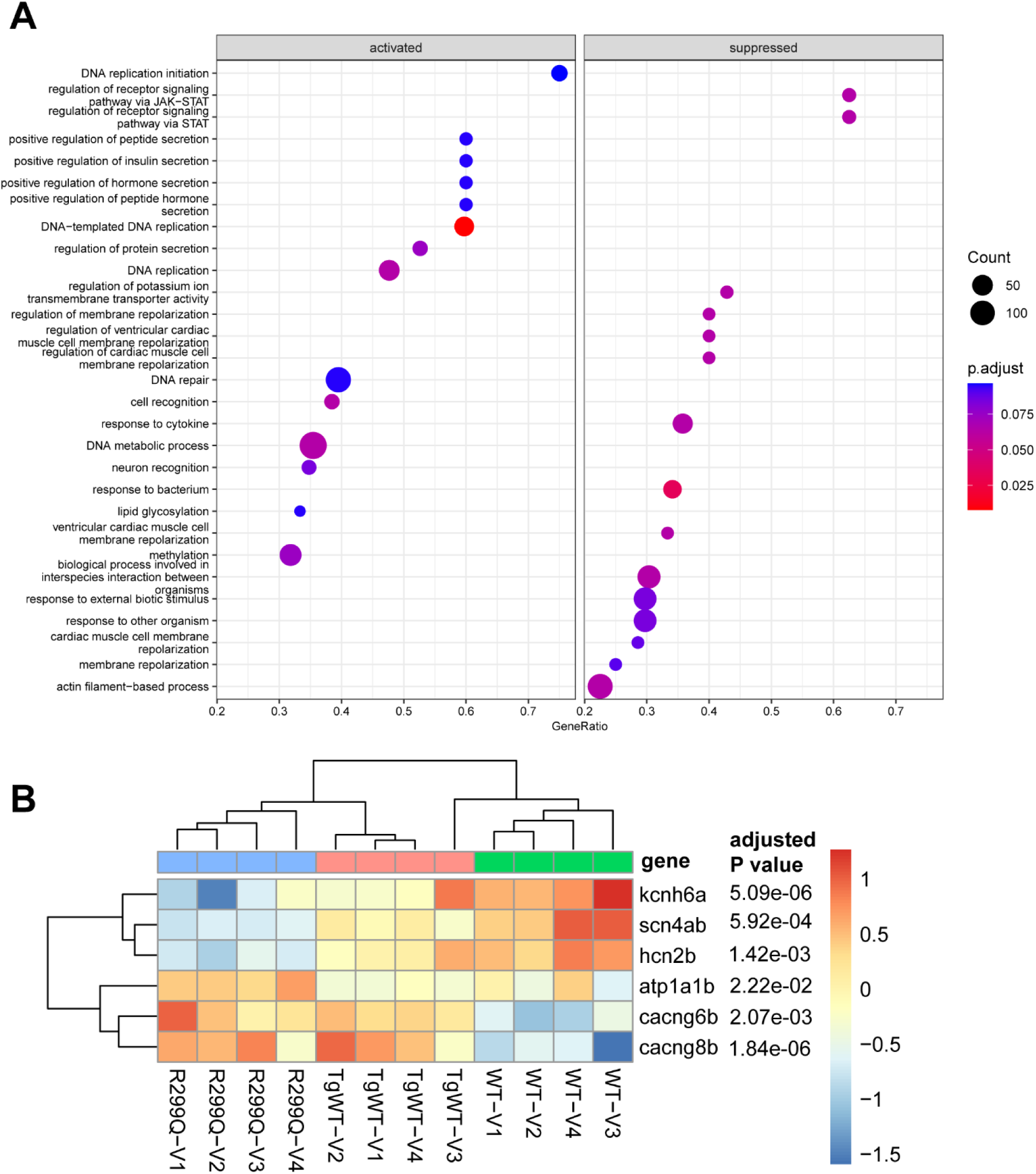
RNA sequencing analysis of adult ventricular tissues. A. Gene set enrichment analysis of differential expression among TuAB, Tg^WT^, and Tg^R299Q^ ventricles. B. Heatmap of selected significantly differentially modulated genes encoding ion channels and transporters.

**Supplement Figure 5.**
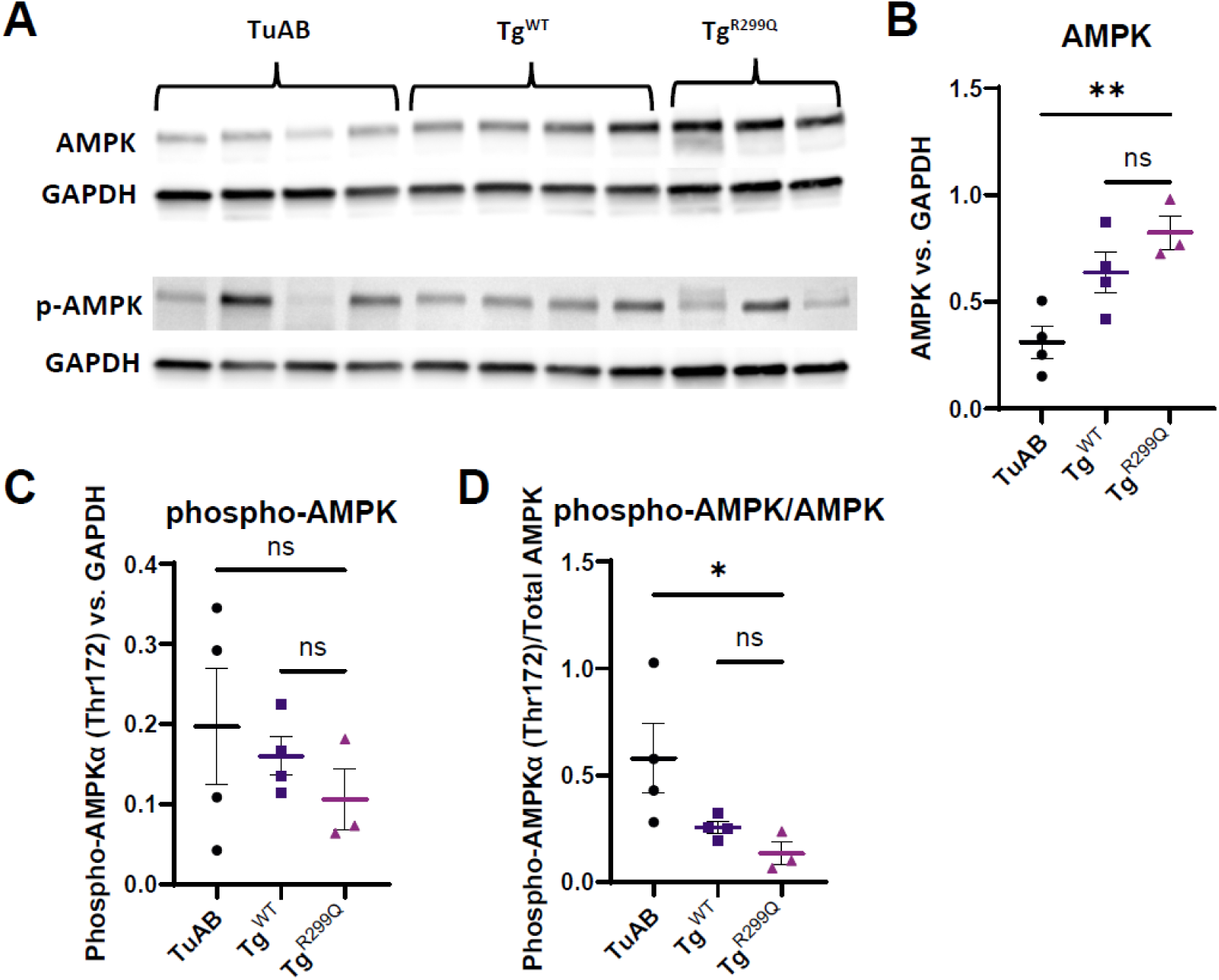
AMPK phosphorylation is down-regulated in Tg^R299Q^ during cardiogenesis. A. Western blot analysis with antibodies for AMPK and phosphorylated AMPK at Thr^172^ of protein lysates from TuAB, Tg^WT^, and Tg^R299Q^ adult hearts. Each loaded sample contains lysates from an individual heart. B-C. Normalized protein expression of AMPK (D) and phospho-AMPK (E) relative to GAPDH among TuAB, Tg^WT^, and Tg^R299Q^. D. The calculated ratio of the normalized phospho-AMPK over total AMPK in TuAB, Tg^WT^, and Tg^R299Q^ adult heart lysates.

**Supplement Figure 6.**
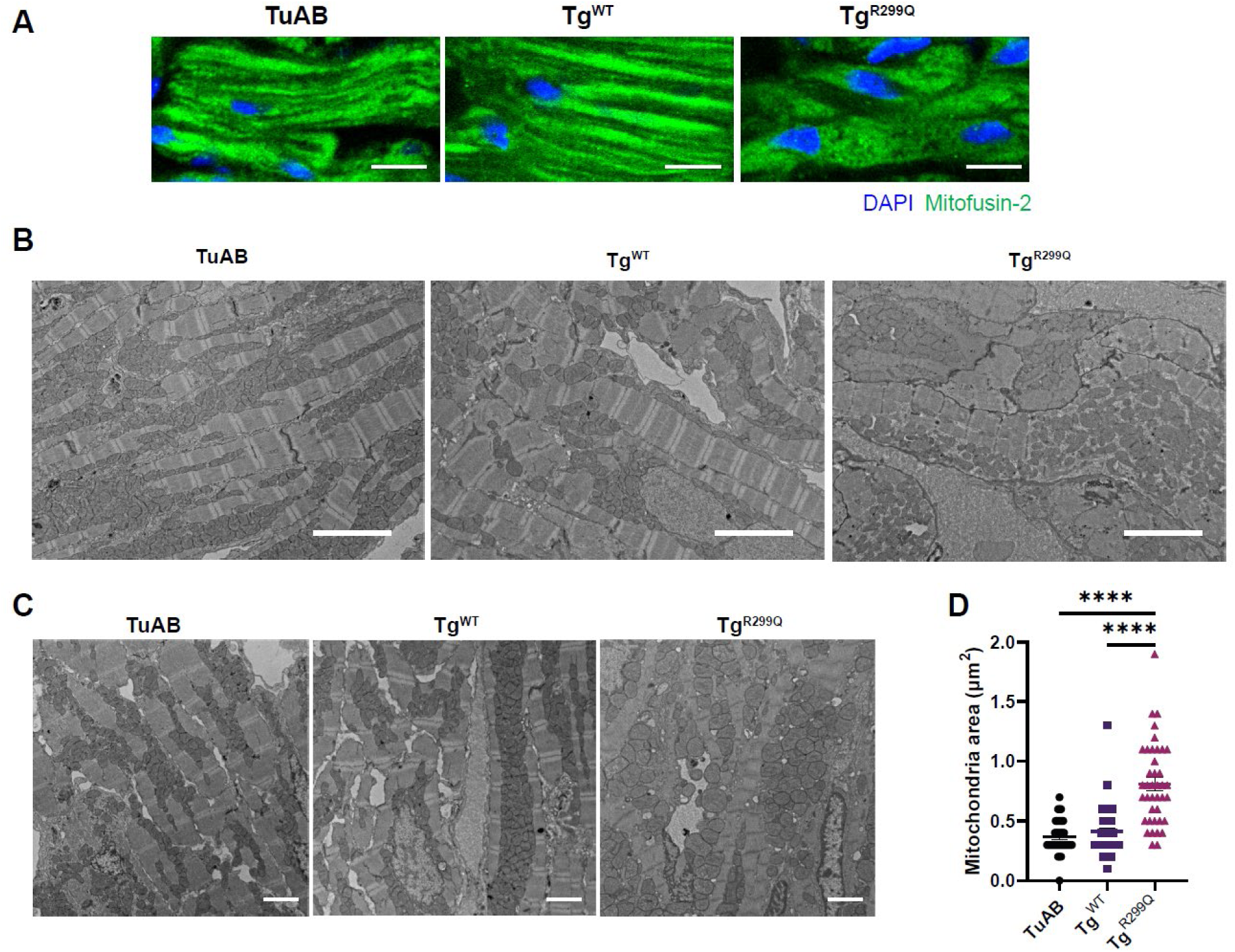
R299Q variant alters the cytoarchitecture of cardiomyocytes. A, Representative immunofluorescence images of ventricular sections stained with Mitofusin-2 (green) and DAPI (blue) from TuAB, Tg^WT^, and Tg^R299Q^ adult ventricles. The scale bars shown are 10 μm. B. TEM images showing cardiomyocyte cytoarchitecture of ventricular sections from TuAB, Tg^WT^, and Tg^R299Q^ adult hearts. The scale bars shown are 4000 nm. C. Example TEM images of ventricular sections of adult hearts from TuAB, Tg^WT^, and Tg^R299Q^ adult hearts. Scale bars indicate 2000 nm. D. Measured mitochondria area from TEM images of TuAB, Tg^WT^, and Tg^R299Q^ ventricular longitudinal sections. Each data point in D indicates individual mitochondria, and four hearts from each genotype were used in this analysis. Results are presented as mean ± S.E.M. **** indicates P < 0.0001.

**Supplement Figure 7.**
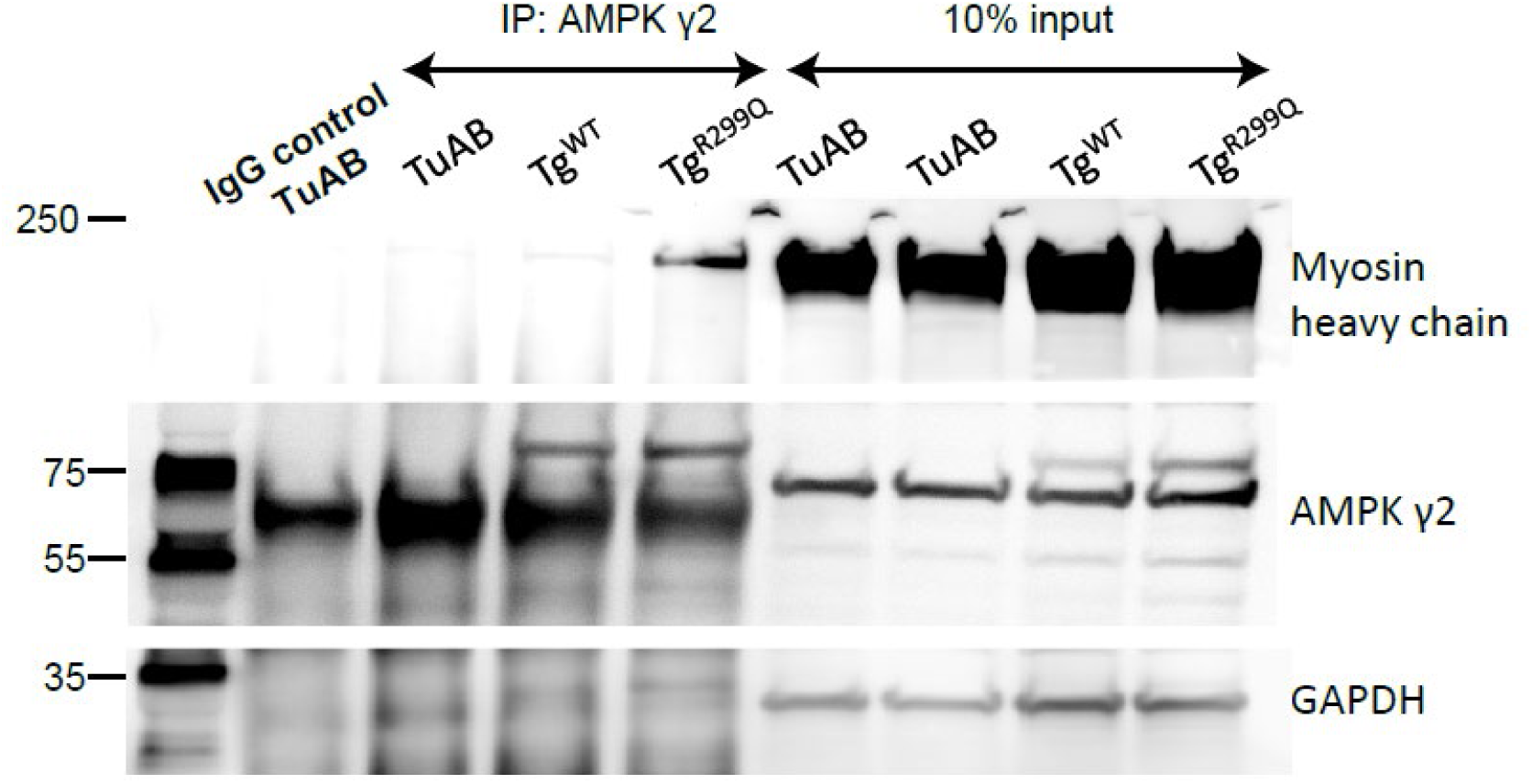
Co-immunoprecipitation of AMPKγ2 and myosin heavy chain. Western blot analysis of immunoprecipitated samples using IgG control beads or AMPKγ2 antibody-conjugated beads (left), along with 10% input from each protein lysate (right). Each lysate was prepared by pooling six adult hearts from TuAB, Tg^WT^, or Tg^R299Q^ fish. Immunoblots for myosin heavy chain, AMPKγ2, and GAPDH are shown.

**Supplement Figure 8.**
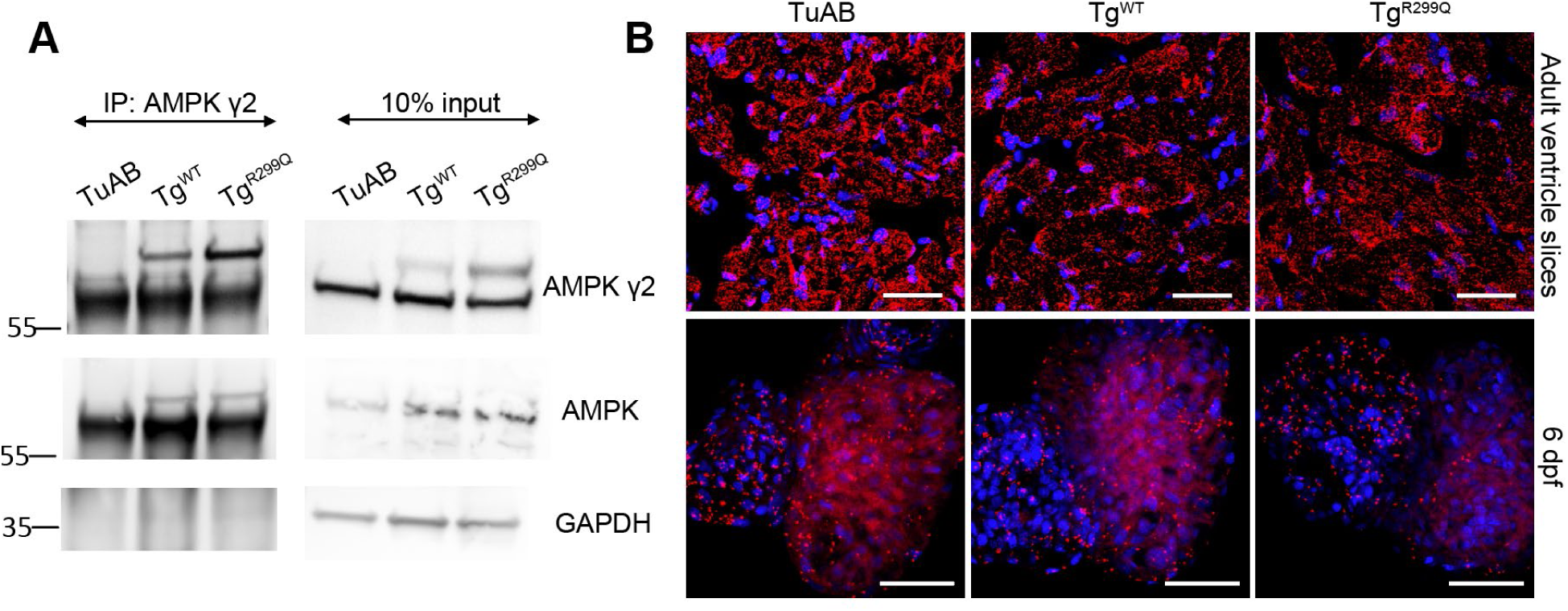
The interaction between AMPK α and γ2 subunits was not affected by the R299Q variant. A. Western blot analysis of immunoprecipitated samples with the AMPKγ2 antibody-conjugated beads (left), and 10% input from each protein lysate (right). Each lysate was prepared by combining six adult hearts dissected from TuAB, Tg^WT^, and Tg^R299Q^ fish respectively. Blots for AMPKγ2, AMPK α, and GAPDH are shown. B-C. Proximity ligation assays (PLAs) using antibodies for AMPKγ2 (rabbit) and AMPK α (mouse). Shown are representative images from ventricular sections of TuAB, Tg^WT^, and Tg^R299Q^ adult hearts (top), and whole hearts isolated from 6 dpf TuAB, Tg^WT^, and Tg^R299Q^ larvae (bottom). Scale bars shown indicate 25 µm.

